# GLP-1(9-36) mediates the glucagonostatic effect of GLP-1 by promiscuous activation of the glucagon receptor

**DOI:** 10.1101/785667

**Authors:** Claudia Guida, Caroline Miranda, Ingrid Wernstedt Asterholm, Davide Basco, Anna Benrick, Belen Chanclon, Margarita V. Chibalina, Matthew Harris, Joely Kellard, Laura J. McCulloch, Joana Real, Nils J.G. Rorsman, Ho Yan Yeung, Frank Reimann, Makoto Shigeto, Anne Clark, Bernard Thorens, Patrik Rorsman, Graham Ladds, Reshma Ramracheya

## Abstract

The incretin hormone glucagon-like peptide 1(7-36) (GLP-1(7-36)) stimulates insulin and inhibits glucagon secretion. The mechanisms by which GLP-1 suppresses glucagon release are unclear as glucagon-secreting α-cells express GLP-1 receptors (GLP-1Rs) at very low levels. Here, we examine the underlying mechanisms. We find that both GLP-1(7-36) and its degradation product GLP-1(9-36) inhibit glucagon secretion at physiological (pM) concentrations. Whereas the effect of GLP-1(7-36) is sensitive to PKA inhibition, GLP-1(9-36) exerts its effect by a PKA-independent mechanism sensitive to pretreatment with pertussis. The glucagonostatic effects of both GLP-1(7-36) and (9-36) are retained in islets from *Glp1r* knockout mice but only GLP-1(9-36) remains glucagonostatic in the presence of the DPP-4 (the peptidase catalyzing the formation of GLP-1(9-36)) inhibitor sitagliptin. Glucagon receptor (GCGR) antagonism specifically prevents the inhibitory effects of GLP-1(9-36) whilst not affecting that of GLP-1(7-36). We conclude that GLP-1(7-36) and GLP-1(9-36) regulate glucagon secretion via interaction with GLP-1R and GCGR, respectively.

**Highlights:**

- GLP-1(7-36) and GLP-1(9-36) inhibit glucagon secretion from alpha-cells
- GLP-1(7-36) and (9-36) retain glucagonostatic effect in Glp1r^-/-^ islets
- GLP-1(7-36) and (9-36) activate distinct signal transduction mechanisms
- GLP-1(7-36) acts via GLP-1R and GLP-1(9-36) via GCGR

## Introduction

Glucagon is the body’s principal hyperglycemic hormone (Cryer, 2015). In both type 1 (T1D) and type 2 diabetes (T2D), hyperglycemia results from a combination of insufficient insulin secretion and oversecretion of glucagon (Unger and Orci, 1975). Whereas the insulin secretion defect has attracted much attention, the dysregulation of glucagon secretion in diabetes remains, by comparison, an understudied area.

The incretin hormone glucagon-like peptide 1 (GLP-1) is secreted by the L-cells of the gut as GLP-1(7-36) which exerts a strong hypoglycemic effect by stimulating insulin secretion in the β-cells (Holst, 2007). Following its release, GLP-1(7-36) is quickly degraded by dipeptidyl peptidase 4 (DPP-4) to form the metabolite GLP-1(9-36) (Baggio and Drucker, 2007). We will refer to the two peptides by their full names and reserve the use of ‘GLP-1’ to describe total GLP-1. Plasma GLP-1 levels range between 10-50 pM of which only <20% is GLP-1(7-36) and the rest exists as GLP-1(9-36) (Orskov et al., 1994).

GLP-1(7-36) exerts a glucagonostatic effect (i.e. inhibits glucagon secretion in the pancreatic α-cells). Although this effect is thought to account for ∼50% of the peptide’s hypoglycemic action (Hare et al., 2010), the cellular mechanisms underlying the glucagonostatic effect are poorly understood compared to the insulinotropic effects, which are fairly well characterized (Gromada et al., 1998).

Most studies indicate that GLP-1 receptors (encoded by *Glp1r*) are expressed at very low levels (if at all) in glucagon-secreting α-cells (De Marinis et al., 2010; Nakashima et al., 2018; Ramracheya et al., 2018; Richards et al., 2014) (but see (Zhang et al., 2019)). It was therefore argued that GLP-1(7-36) exerts its inhibitory effect on glucagon secretion by a paracrine mechanism mediated by factor(s) secreted by the neighboring β- and δ-cells (e.g. insulin and somatostatin, respectively) within the pancreatic islets (Orgaard and Holst, 2017). Yet, GLP-1(7-36) robustly inhibit glucagon secretion in isolated islets when tested at non-physiological nanomolar levels by mechanisms that cannot be accounted for by paracrine signals (De Marinis et al., 2010; Ramracheya et al., 2018). We have previously proposed that this effect is mediated by activation of the limited number of GLP-1 receptors present in α-cells and that this, via a small increase in cAMP, results in inhibition of the voltage-gated Ca^2+^ channels linked to exocytosis of glucagon-containing secretory granules.

We have now explored the role of the ‘classical’ GLP-1 receptor in GLP-1-induced effects on glucagon release by extending our studies to *Glp1r* knockout mice (Scrocchi et al., 1996). The data suggest that activation of GLP-1Rs only mediate part of the glucagonostatic effect of GLP-1 and that GLP-1 also exerts *Glp1r*- and PKA-independent effects. Here we have also studied the effects of GLP-1(7-36) and its metabolite, GLP-1(9-36) on human islet function, and explored the possible involvement of alternative receptors and the intracellular signaling mechanisms involved.

## Results

### Dose-dependent inhibition of glucagon secretion by GLP-1(7-36) and GLP-1(9-36)

We measured glucagon secretion in isolated mouse and human pancreatic islets exposed to 1 mM glucose (this glucose concentration represents a strong stimulus of glucagon secretion) and increasing concentrations of GLP-1(7-36) (Figure 1A-B). In both mouse and human islets, GLP-1(7-36) produced a concentration-dependent inhibition of glucagon secretion that was maximal at 1-10 pM; high concentrations (0.1-100 nM) of GLP-1(7-36) were less efficacious than picomolar levels. In the remainder of the manuscript, GLP-1(7-36) was therefore tested at 1 or 10 pM.

**Figure 1:**
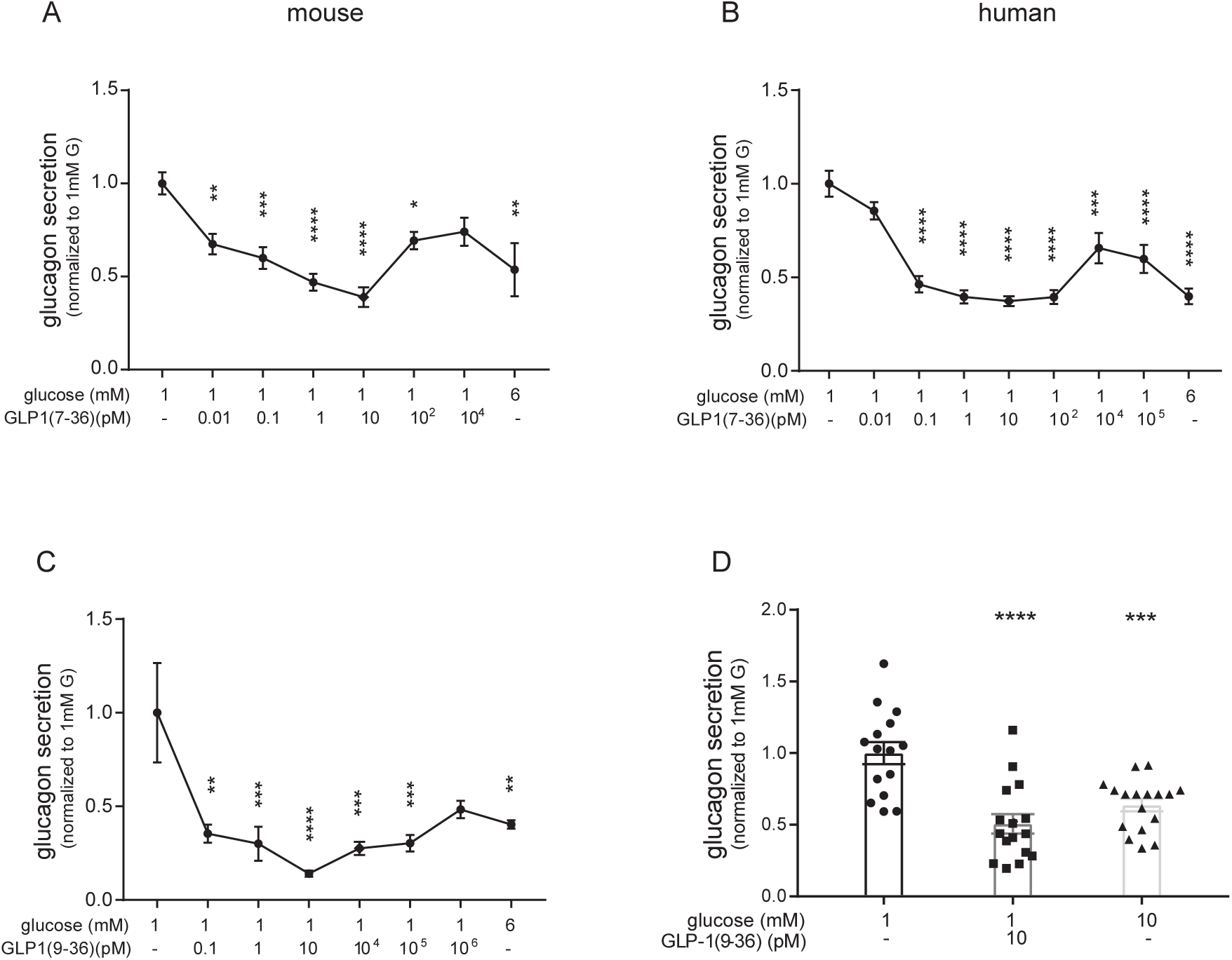
GLP-1 inhibits glucagon secretion. (**A-B**) Effects of increasing concentrations of GLP-1(7-36) on glucagon secretion in isolated mouse (A; n=8-11 using islets from 10 mice: glucagon secretion at 1 mM glucose average 3.8±0.2 pg/islet/h) and human (B; n=5-15 using islets from 4 donors, glucagon secretion at 1 mM glucose averaged 2.2±0.4 pg/islet/h) pancreatic islets. (**C-D**) As in (A and B) but using the metabolite GLP-1(9-36). Glucagon secretion was normalized to that at 1 mM glucose (in C: 2.6±0.2 pg/islet/h, n=4 using islets from 4 mice; in D: 9.2±1 pg/islet/h, n=15-17 using islets from 4 donors). *P<0.05, **P<0.01, ***P<0.001, ****P<0.0001 versus 1mM glucose; 1-way ANOVA with Dunnett’s post-hoc test.

Pancreatic islets are extensively vascularized and the disruption of blood flow during the islet isolation process may lead to intra-islet accumulation of secreted products to unphysiological levels. In static incubations of mouse islets, the glucagon and insulin concentrations measured at the end of the 1-h incubations averaged 69±5 pM (n=8) and 130±13 pM (n=8), respectively. To ascertain that intra-islet paracrine factors operating under static incubation conditions do not interfere with the responses, we also tested GLP-1(7-36) at 1 pM and 10 nM using the dynamic perfused mouse pancreas paradigm. In these experiments, the pancreas was perfused at the physiological rate (∼0.3 ml/min) and in normal direction of vascular flow (51). This methodology has the additional advantage that the islets have not been subjected to any mechanical or enzymatic treatment. When tested at 1 mM glucose, application of 1 pM and 10 nM GLP-1(7-36) inhibited glucagon secretion by 61±8% (n=3) and 40±3.5% (n=3), respectively (Figure S1A-B). These results are in good agreement with those obtained in the static incubation setting, thus validating the latter experimental approach.

In both mouse and human islets, the maximum inhibitory effect of GLP-1(7-36) was comparable to that produced by 6 mM glucose (the glucose concentration with the strongest glucagonostatic effect in both mouse and human islets (Walker et al., 2011). When tested at supraphysiological concentrations (10 nM), GLP-1(7-36) augments the inhibitory effect of high glucose on glucagon secretion in both mouse (De Marinis et al., 2010) and human islets (Ramracheya et al., 2018). However, the effects of physiological concentrations of GLP-1(7-36) (10 pM) and glucose were not additive in isolated human islets (Figure S1C), possibly because a low concentration of GLP-1(7-36) exerts a stronger glucagonostatic effect on its own.

Following its release, GLP-1(7-36) is quickly degraded by dipeptidyl peptidase 4 (DPP-4) to form the metabolite GLP-1(9-36) (Baggio and Drucker, 2007). In mouse islets, GLP-1(9-36) inhibited glucagon secretion as potently as GLP-1(7-36) (Figure 1C); the greatest inhibitory effect was produced by 10 pM. As was the case with GLP-1(7-36), high concentrations (≥1 nM) of GLP-1(9-36) were less efficacious than low concentrations (1-10 pM). The maximum inhibitory effect of GLP-1(9-36) (80% at 1 pM) in fact exceeded that produced by 6 mM glucose. In human islets, GLP-1(9-36) was also strongly inhibitory and reduced glucagon secretion at 1 mM glucose by 50% when tested at a concentration of 10 pM (Figure 1D), similar to that produced by 10 mM glucose (50%).

We excluded the possibility that GLP-1(9-36) inhibits glucagon secretion by a paracrine effect mediated by stimulation of insulin or somatostatin secretion (Figure S2A-D).

Collectively, these data therefore suggest that glucagon secretion is under strong tonic inhibition by circulating levels of GLP-1 (10-50 pM) and that these effects are mediated by direct effects on the α-cells.

### Glucagonostatic and insulinotropic effects of GLP-1 are mediated by pharmacologically distinct mechanisms

The insulinotropic effects of GLP-1(7-36) are shared with the GLP-1 receptor agonist exendin-4, which shows only 53% homology to GLP-1(7-36) (Underwood et al., 2010). Exendin-4 (10 pM) potentiated glucose (6 mM)-induced insulin secretion by 100% in mouse and human islets (Figure 2A-B), comparable to that previously observed for GLP-1(7-36) (Shigeto et al., 2015). By contrast, exendin-4 (0.1 pM-100 nM) was without glucagonostatic effects in mouse islets (Figure 2C). Due to limited access, in human islets only a single concentration of exendin-4 (10 pM) was tested on glucagon secretion but the same negative result was observed (Figure 2D).

**Figure 2:**
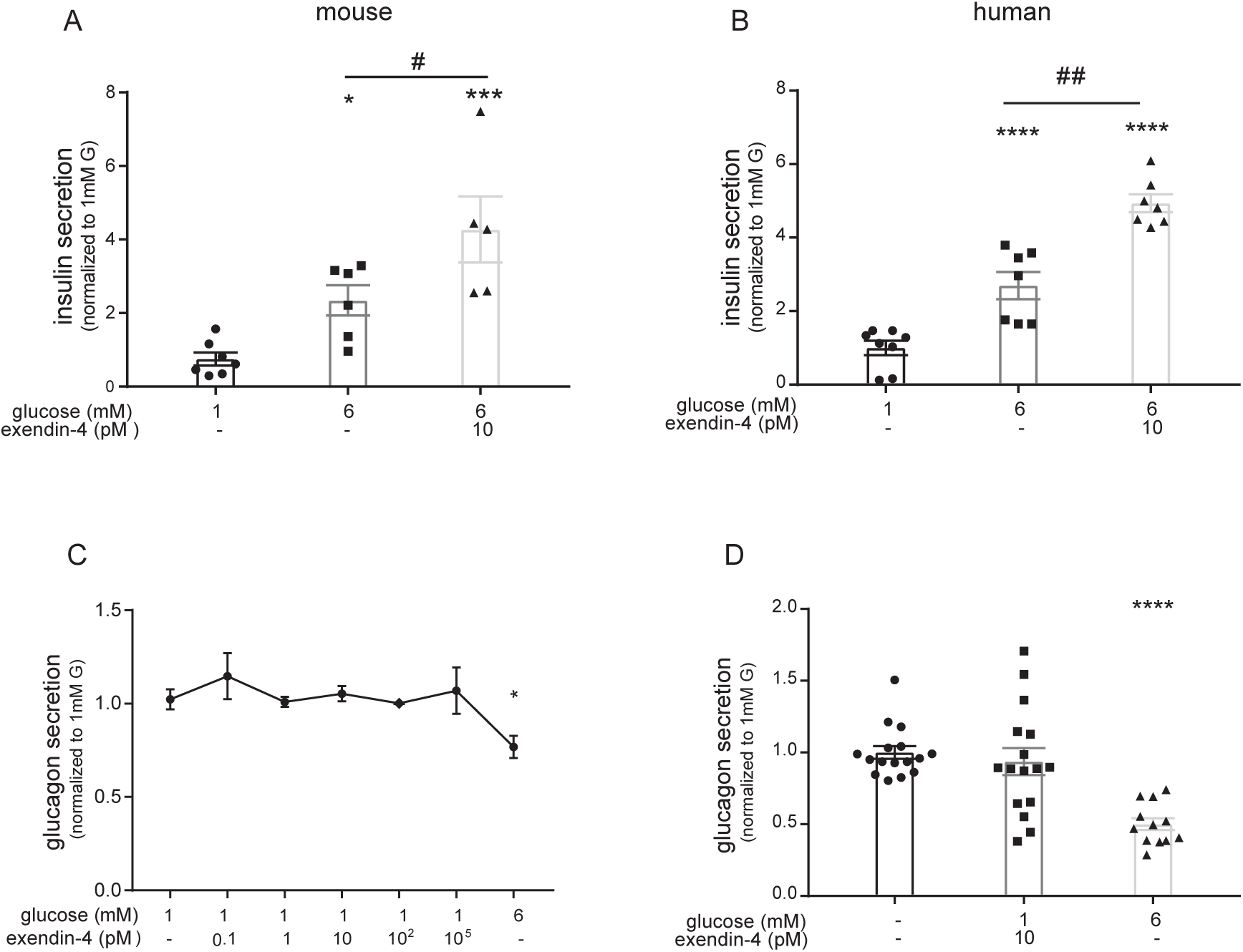
Differential pharmacology of insulinotropic and glucagonostatic effects of GLP-1. (**A-B**) Effects of 10 pM exendin-4 on insulin secretion in mouse. Glucagon secretion was normalized to that at 1 mM glucose (in A: 7.4±1.9 pg/islet/h, n=3 using islets from 3 mice; in B: 6.8±1.9 pg/islet/h, n=7 using islets from 2 donors). (**C-D**) Effects of GLP-1 receptor agonist exendin-4 on glucagon secretion at 1 and 6 mM glucose in mouse and human pancreatic islets. Glucagon secretion was normalized to that at 1 mM glucose (in C: 4.7±0.3 pg/islet/h, n=7-11 using islets from 12 mice; in D: 10.2±1.8 pg/islet/h, n=12-16 using islets from 4 donors). *P<0.05, **P<0.01, ****P<0.0001 versus 1 mM glucose; ## P<0.01 for indicated comparison,1-way ANOVA with Dunnett’s post-hoc test.

### GLP-1(7-36) and GLP-1(9-36) signal by PKA-dependent and -independent mechanisms, respectively

We compared the glucagonostatic effects of physiological concentrations of GLP-1(7-36) and GLP-1(9-36) in the absence and presence of the PKA-inhibitor 8-Br-Rp-cAMPS. Under control conditions, GLP-1(7-36) (10 pM) inhibited glucagon secretion by 48%, which was reduced to 25% in the presence of 8-Br-Rp-cAMPS (10 µM; p<0.01). By contrast, the effect of GLP-1(9-36) (10 pM) was not at all affected by the PKA inhibitor. In this experimental series, the inhibitory effect of GLP-1(9-36) averaged 42% and 40% (p=0.7) in the absence and presence of 8-Br-Rp-cAMPS, respectively (Figure 3B). Thus, it appears that whereas the inhibitory effects of GLP-1(7-36) are partially PKA-dependent, those of GLP-1(9-36) are entirely PKA-independent. Notably, the PKA inhibitor had no effect on glucagon secretion at 1 mM glucose. This argues that cAMP-dependent activation of PKA does not account for the stimulation of glucagon secretion at low glucose, at variance with a recent suggestion (Yu et al., 2019).

**Figure 3:**
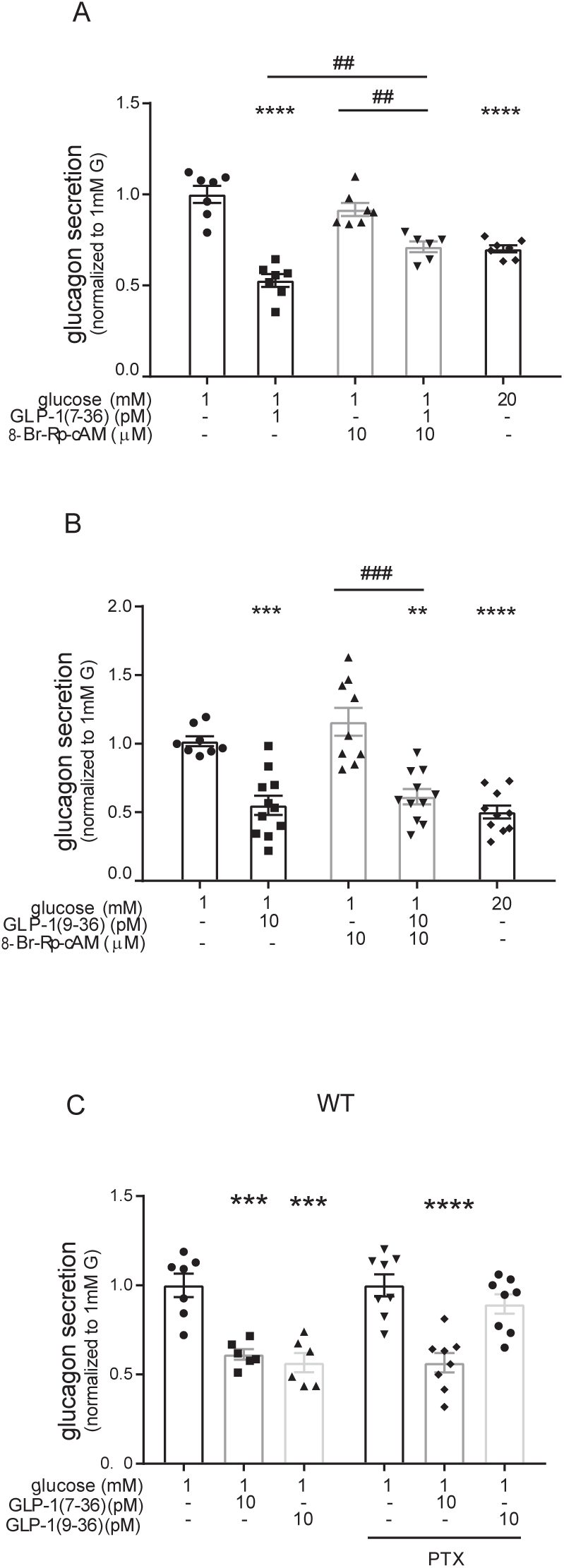
PKA-dependent and G_i_–dependent glucagonostatic effects of GLP-1(7-36) and GLP-1(9-36) (**A**) Effects of GLP-1(7-36) on glucagon secretion in the absence and presence of the PKA-inhibitor 8-Br-Rp-cAMPS as indicated (n=7 using islets from 11 mice, glucagon secretion at 1 mM glucose averaged 2.8±0.13 pg/islet/h). Glucose (G) was included at 1 and 20 mM as indicated. (**B**) As in (A) but using GLP-1(9-36) (n=8-11 using islets from 10 mice, glucagon secretion at 1 mM averaged 1.8±0.3 pg/islet/h, at 1 mM G+8-Br-Rp-cAMP=2.1±0.2 pg/islet/h, ***P<0.001, ****P<0.0001 versus 1mM glucose, ##P<0.01, ###P<0.001 for indicated comparisons; 1-way ANOVA with Dunnett’s post-hoc test. (**C**) Effects of 10 pM GLP-1(7-36) or GLP-1(9-36) on glucagon release in wild type (WT) islets under control conditions and after pre-treatment with pertussis toxin (PTX). Glucagon secretion normalised to that at 1 mM glucose which averaged 3.05±0.70 pg/islet/h and 3.9±0.25 pg/islet/h the absence and presence of PTX (n=6-8 using islets from 11 mice)

### GLP-1(9-36), but not GLP-1(7-36), signals via a G_i_-dependent mechanism

We hypothesized that the PKA-independent effects of GLP-1(9-36) are mediated by activation of an inhibitory GTP-binding protein (G_i_). We next compared the effects of GLP-1(7-36) and (9-36) in islets pretreated with pertussis toxin (PTX) to inhibit G_i_ (Asano et al., 1984). Under control conditions (without PTX pre-treatment), GLP-1(7-36) and (9-36) (both tested at 10 pM) inhibited glucagon secretion by >50%. Following pre-treatment with PTX (100 ng/ml for >4h), the inhibitory effect of GLP-1(9-36) was abolished whereas that of GLP-1(7-36) was unaffected (Figure 3C). These experiments were not repeated in islets from human donors because pretreatment of human islets with PTX exerts a paradoxical strong glucagonostatic effect on its own (Walker et al., 2011).

### The glucagonostatic effect of GLP-1 is retained in Glp1r^-/-^ islets

Collectively, the data presented in Figures 1-3 suggest that GLP-1(7-36) stimulates insulin secretion and inhibits glucagon release by pharmacologically distinct mechanisms that involve separate signal transduction pathways. These findings raise the interesting possibility that the glucagonostatic effects of GLP-1(7-36) and (9-36) involve a receptor distinct from the previously characterized GLP-1 receptor (GLP-1R). We explored this possibility by comparing the effects of GLP-1(7-36) and GLP-1(9-36) on insulin and glucagon secretion in islets isolated from control (*Glp1r*^*+/+*^) and GLP-1 receptor knockout (*Glp1r*^*-/-*^) mice.

In wild-type islets, increasing glucose from 1 to 6 mM stimulated insulin secretion by 1.9-fold. Treatment with GLP-1(7-36) (10 pM) resulted in an additional 1.4-fold increase of insulin secretion. In *Glp1r*^*-/-*^ islets, increasing glucose from 1 to 6 mM also stimulated insulin secretion by 125% but GLP-1(7-36) (10 pM) did not lead to any further stimulation (Figure 4A).

**Figure 4:**
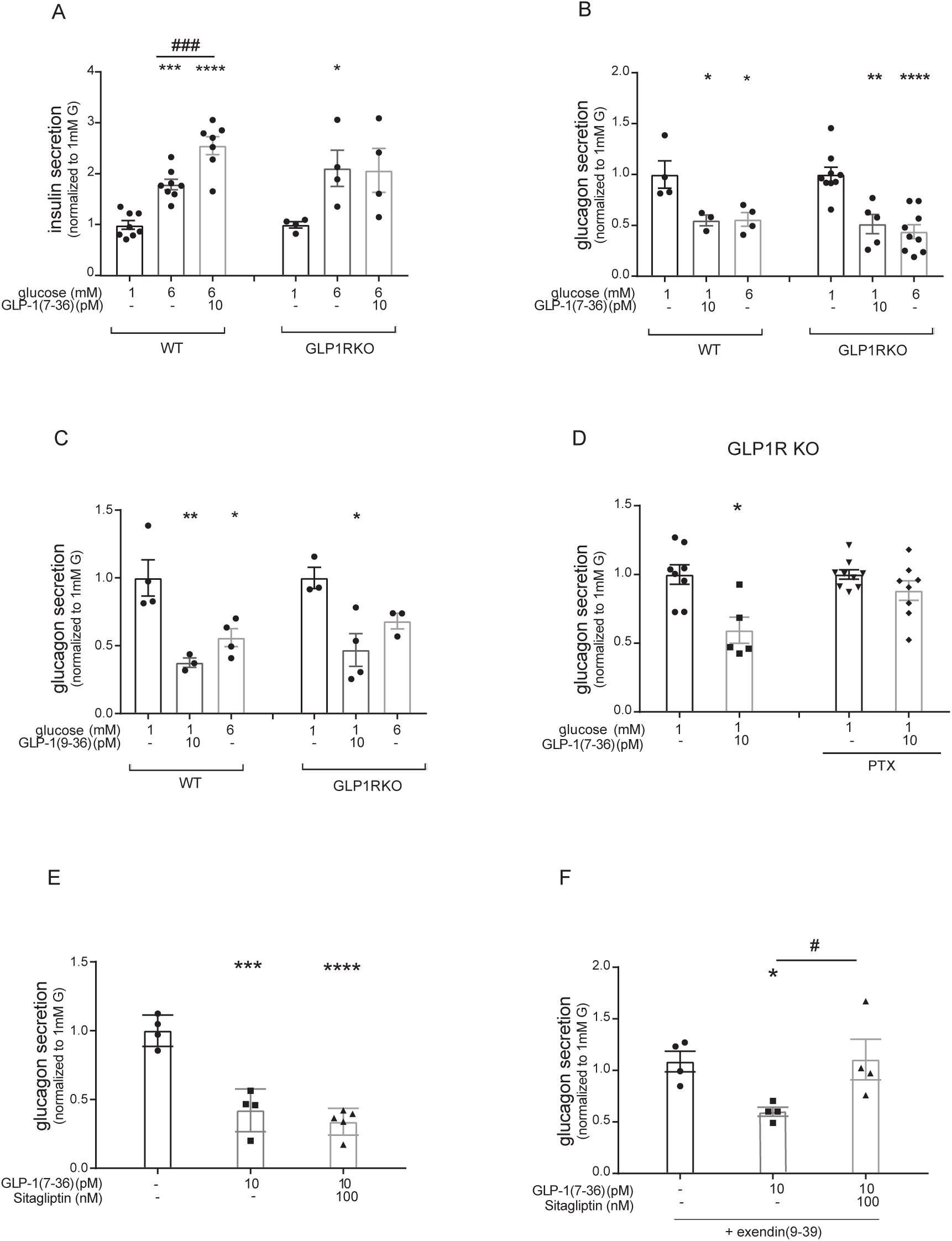
GLP-1(7-36) and GLP-1(9-36) retain glucagonostatic effects in Glp1r-/- mouse islets. **(A)** Effects of 10 pM GLP-1(7-36) on insulin secretion at 1 and 6 mM glucose in islets isolated from wild-type (WT) and *Glp1r*^*-/-*^ (GLP1R-KO) mice (n=4-8 using islets from 4 mice of each genotype; insulin secretion normalised to that at 1 mM glucose). **(B)** Effects of 10 pM GLP-1(7-36) on glucagon secretion in wild-type and *Glp1r*^*-/-*^ islets (n=4-9 using islets from 4 mice of each genotype; glucagon secretion normalised to that at 1 mM glucose which averaged 3.4±0.46 pg/islet/h and 2.6±0.2 pg/islet/h in wildtype and *Glp1r*^*-/-*^ islets, respectively). **(C)** As in (B) but testing effects of 10 pM GLP-1(9-36) (n=3-4 using islets from 3 mice of each genotype; glucagon secretion normalised to that at 1 mM glucose which averaged 3.5±0.5 pg/islet/h and 2.2±0.17 pg/islet/h in wildtype and *Glp1r*^*-/-*^ islets, respectively. (**D**) Effects of 10 pM GLP-1(7-36) or GLP-1(9-36) on glucagon release in islets from *Glp1r*^*-/-*^ mice. Glucagon secretion normalised to that at 1 mM glucose which averaged 4.3±0.6 pg/islet/h and 5.7±0.95 pg/islet/h in the absence and presence of PTX (n=6-9 using islets from 10 mice). (**E-F**) Effects of 10 pM GLP-1(7-36) on glucagon secretion in the presence of the DPP-4 inhibitor sitagliptin (100 nM) alone (E, glucagon secretion at 1mM G=4.07±0.2 pg/islet/h) and with the GLP-1 receptor antagonist exendin(9-39) (1 μM) (F, glucagon secretion at 1 mM G=2.3±0.26 pg/islet/h) (n=4 using islets from 4 mice). *P<0.05, **P<0.01, ***P<0.001, ****P<0.0001 versus 1mM glucose; # P<0.05 for indicated comparison; 1-way ANOVA with Dunnett’s post-hoc test.

The effects of GLP-1(7-36) on glucagon secretion were tested at 1 mM glucose. In control islets, GLP-1(7-36) (10 pM) inhibited glucagon secretion by ∼50%. The inhibitory effects of GLP-1(7-36) persisted in *Glp1r*^*-/-*^ islets where it amounted to a 55% reduction in glucagon release (Figure 4B). Similarly, GLP-1(9-36) (10 pM) was equally inhibitory in *Glp1r*^*-/-*^ and control islets, suppressing glucagon secretion by ∼60% (Figure 4C). The inhibitory effects of both, GLP-1(7-36) and GLP-1(9-36) on glucagon secretion were comparable to the suppression resulting from increasing the glucose concentration from 1 to 6 mM. The reduction of glucagon secretion at high glucose concentration was not affected by ablation of *Glp1r* (Figure 4B-C).

We point out that the mouse model used for these experiments is a general knockout and the possibility that GLP-1 exerts its inhibitory action by a paracrine effect by activation of GLP-1Rs in δ- and β-cells can therefore be excluded. Thus, whereas the insulinotropic effect of GLP-1 is mediated by the previously characterized (cloned) GLP-1R, the glucagonostatic effects are probably mediated by activation of different receptor(s).

We next tested the capacity of GLP-1(7-36) to suppress glucagon secretion in islets from *Glp1r*^*-/-*^ mice following inactivation of G_i_. In keeping with the data in Figure 4B, GLP-1(7-36) retained its capacity to inhibit glucagon secretion in *Glp1r*-deficient islets. However, unlike what was observed in control (wild-type) islets, GLP-1(7-36) did not inhibit glucagon secretion in PTX-pretreated *Glp1r*^*-/-*^ islets (Figure 4D).

The latter finding raises the interesting possibility that the *Glp1r*-independent inhibitory effects of GLP-1(7-36) on glucagon secretion are actually mediated by its breakdown product GLP-1(9-36). DPP-4, the enzyme catalyzing this reaction, is expressed and functionally-active in isolated pancreatic islets (Guida et al., 2017). To explore this hypothesis, we tested the effects of GLP-1(7-36) (10 pM) in the presence of the DPP-4 inhibitor sitagliptin (100 nM) (Deacon, 2018) and compared the effects on glucagon secretion in the absence or presence of the GLP-1 receptor antagonist exendin(9-39) (100 nM) (Figure 4E-F). Consistent with the data in *Glp1r*-deficient islets (Figure 4B), GLP-1(7-36) retained an inhibitory effect on glucagon release in the presence of exendin(9-39). However, co-application of sitagliptin and exendin(9-39) abolished the inhibitory effect of GLP-1(7-36) (Figure 4F). Importantly, sitagliptin was without stimulatory effect on glucagon secretion when tested in the absence of exendin(9-39) (Figure 4E).

### GLP-1(9-36) activates the glucagon receptor

We compared expression of *Glp1r* and other receptors that can influence glucagon secretion. Consistent with previous reports(De Marinis et al., 2010), expression of *Glp1r* and *Gcgr* in mouse α-cells is lower than that in β-cells. Notably, *Glp1r, Gipr* and *Gpr119* are all expressed in α-cells at levels comparable to *Adrb1* (the β_1_-adrenergic receptor) (Figure S3A-B). Activation of β-adrenergic receptors has been shown to exert a dual effect on glucagon secretion; whereas low concentrations inhibit glucagon secretion by a PKA-dependent effect, high concentrations stimulate glucagon release by activation of Epac2. Similar dual and dose-dependent effects on glucagon secretion are produced by the adenylate cyclase activator forskolin(De Marinis et al., 2010). Analyses of published RNAseq data of human islet cells (Blodgett et al., 2015) indicated that whereas *GLP1R* and *GCGR* are expressed at low levels in α-cells (0.1-5% of that in β-cells), it is greater than what can be accounted for by possible contamination by β-cells. By contrast, *GIPR, ADRB1* and *GPR119* were expressed at >10- to 100-fold higher levels (Figure S3C-D). Notably, both *ADRB1* and *GPR119* are expressed at much higher levels in α-cells than in β-cells.

We speculated that the *Glp1r*-independent effects of GLP-1(7-36) and GLP-1(9-36) might be mediated by activation of the GCGR. In HEK 293 cells expressing human *GCGR*, glucagon increased cAMP production with an EC_50_ of ∼1 nM, an effect that was antagonized by the glucagon receptor antagonist L-168049 (Cascieri et al., 1999) (Figure 5A). Intriguingly, GLP-1(9-36) also activated the glucagon receptor and increased cAMP content with an EC_50_ of 30 nM (Figure 5B). This effect was also antagonized by L-168049, confirming that the action is due to activation of the glucagon receptor (Figure 5B). While the potency is relatively low in these cells, this is probably due to the insensitive nature of the cAMP reporter assay used.

**Figure 5:**
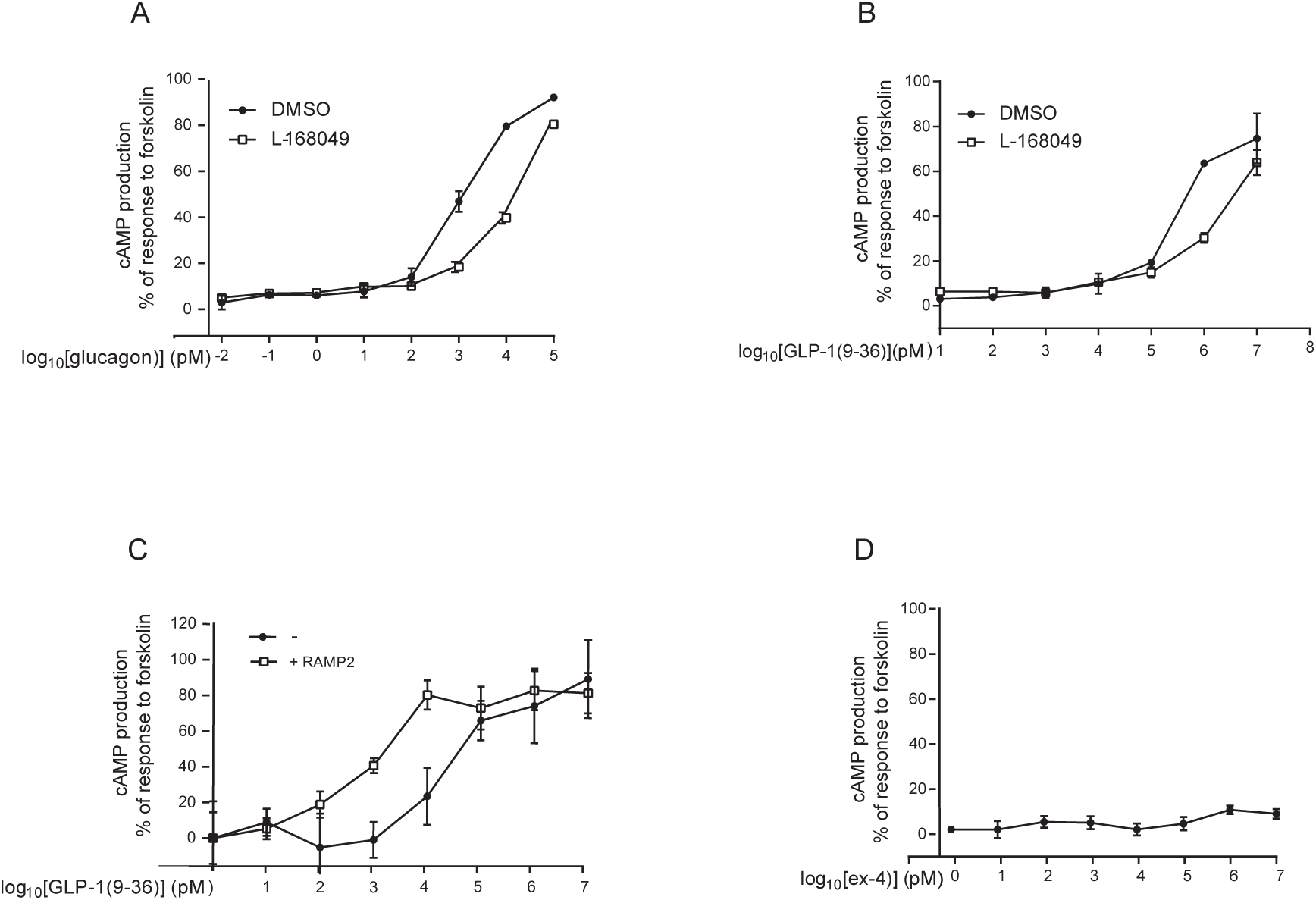
GLP-1(9-36) activates the glucagon receptor. (**A**) cAMP accumulation was determined in [ΔCTR] HEK 293 cells transiently transfected with the GCGR and stimulated for 15 min with glucagon (n=6) (**B**) cAMP accumulation was determined in [ΔCTR] HEK 293 cells transiently transfected with the GCGR (-) or with GCGR +RAMP2 and stimulated for 15 min with GLP-1(9-36) (n=4) Data expressed as percentage of cAMP production relative to GCGR alone. (**C**) Effects of GLP-1(9-36) on cAMP production in the absence and presence of L-168049 as indicated (n=10) (**D**) cAMP accumulation was determined in [ΔCTR] HEK 293 cells transiently transfected with the GCGR and stimulated for 15 min with exendin-4 (n=6). All data, except panel C, are expressed as percentage of cAMP production measured in the presence of 100 μM forskolin.

Receptor activity-modifying proteins (RAMPs) modulate the pharmacology of G protein-coupled receptors (McLatchie et al., 1998). In human islets, RAMP2 is expressed at much higher (∼20-fold) levels in the α-than in β-cells (Blodgett et al., 2015) (Figure S3B). Whereas the capacity of GLP-1(7-36) to activate the human GCGR is abolished when the receptor is co-expressed with RAMP2 (Weston et al., 2015), the activating potency of GLP-1(9-36) was enhanced 30-fold; the EC_50_ was reduced from 10 nM to 0.3 nM (Figure 5C). Significantly, and in contrast to both GLP-1(7-36) and (9-36), exendin-4 was without effect on cAMP production in GCGR-expressing HEK 293 cells (Figure 5D).

Given the high expression of *Gpr119*/*GPR119* in mouse and human α-cells, we explored the possibility that activation of this receptor may contribute to GLP-1’s glucagonostatic effects. The Gpr119 agonist AS1269574 inhibited glucagon secretion in isolated mouse islets when applied at concentrations of 0.3-3 nM but was ineffective at higher concentrations (Figure S4A). However, both GLP-1(7-36) and (9-36) remained capable of inhibiting glucagon secretion in islets from *Gpr119*^*-/-*^ mice (Figure S4B-C) and neither GLP-(7-36) nor (9-36) activated GPR119 expressed in HEK 293 cells even when tested at concentrations as high as 10 μM (Figure S4D). We finally excluded the possibility that the GLP-1R agonists mediate their actions through the activation of GIPRs (Figure S5).

### Loss of glucagonostatic effect of GLP-1(7-36) and (9-36) in *Gcgr*^*-/-*^ islets

The receptor activation studies raise the interesting possibility that GLP-1(9-36) may act by activating the glucagon receptors. GCGR is pleiotropically coupled, i.e. activating not only stimulatory G_s_ but also the inhibitory G_i_ GTP-binding proteins (Weston et al., 2015; Weston et al., 2014; Wootten et al., 2016). In HEK 293 cells expressing the GCGR, pretreatment with pertussis toxin increased cAMP accumulation in response to GLP-1(9-36) (Figure S6), indicating that GLP-1(9-36) can activate G_i_. It was ascertained that there was no increase in cAMP in HEK 293 cells not expressing GCGR.

The studies with recombinant glucagon receptors and glucagon receptor antagonists suggest that GLP-1(9-36) may activate glucagon receptors (GCGRs). We therefore compared the effects of GLP-1(7-36) (10 pM) and (9-36) (10 pM) in islets from control and *Gcgr*^*-/-*^ mice (Gelling et al., 2003). In wildtype islets, GLP-1(7-36) and GLP-1(9-36) applied at 1 mM glucose inhibited glucagon secretion as strongly as increasing glucose from 1 to 6 mM (Figure 6A).

**Figure 6:**
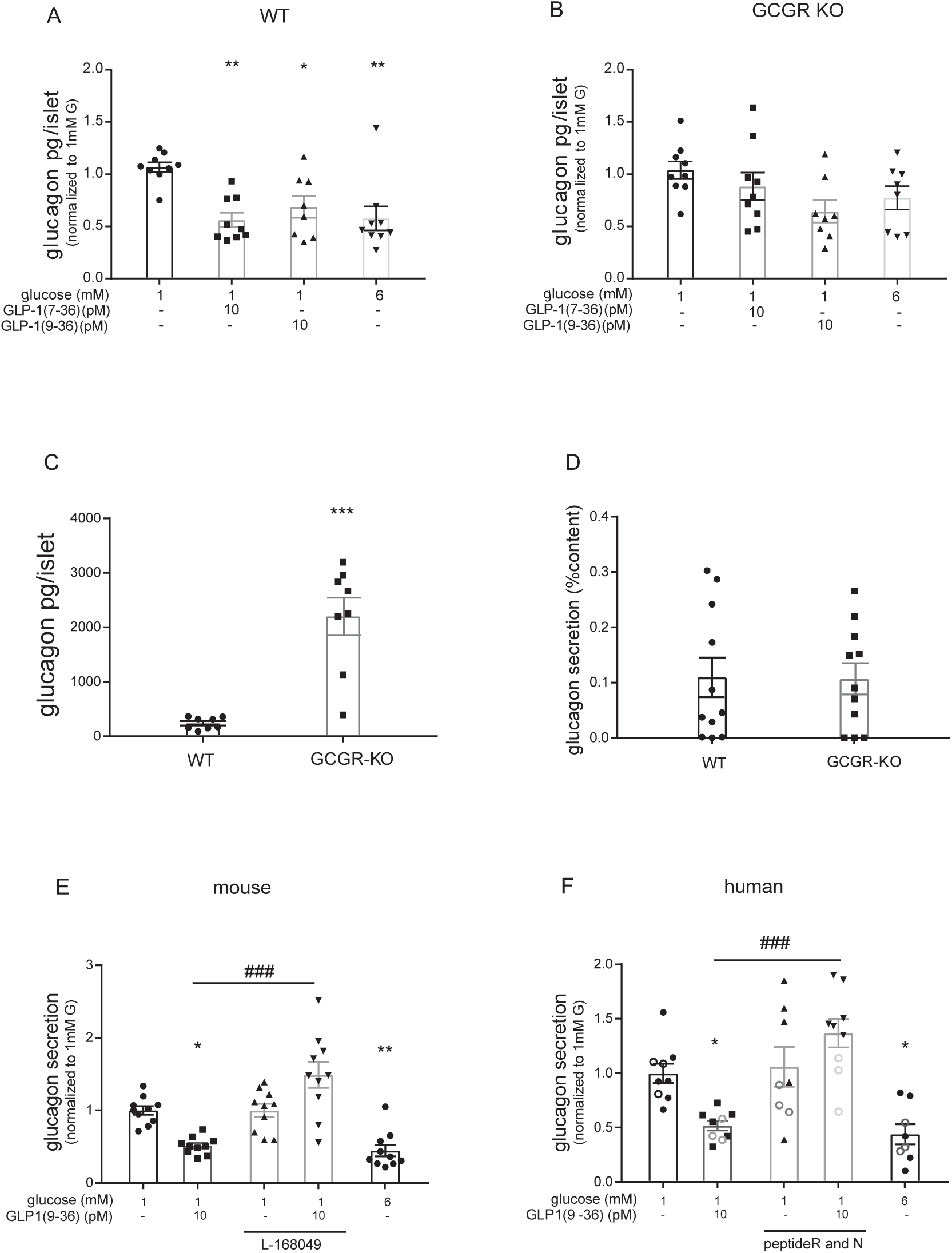
No glucagonostatic effect of GLP-1(9-36) after genetic or pharmacological inhibition of GCGR. (**A-B**) Effects of 10 pM GLP-1(7-36) or 10 pM GLP-1(9-36) on glucagon secretion in islets from wild type (WT; A) and *Gcgr*^*-/-*^ mice (GCGR KO; B). Glucagon secretion has been normalised to that at 1 mM glucose, which averaged 0.83±0.2 pg/islet/h and 4.35±0.76 pg/islet/h in wildtype and *Gcgr*^*-/-*^ islets, respectively (n=9 using islets from 12 mice of each genotype). (**C-D**) glucagon content (C) and glucagon secretion expressed as %content (D) in islets isolated from wild type and Gcgr-/- mice (n=8). **(E)** Effects of 10 pM GLP-1(9-36) on glucagon secretion in the absence or presence of L-168049 (A; n=10 using islets from 7 mice; glucagon secretion at 1 mM glucose averaged 5.9±0.3 pg/islet/h). **(F)** Effect of 10 pM GLP-1(9-36) on glucagon secretion in human islets in absence and presence of 100 nM Peptides ‘R’ or ‘N’ (n=8-9 from 3 donors; glucagon secretion at 1 mM glucose averaged 7.2 ±0.7 pg/islet/h). *P<0.05, **P<0.01,***P<0.001, versus 1mM glucose; ### P<0.001 for indicated comparison, 1-way ANOVA with Dunnett’s post-hoc test.

Consistent with earlier reports (Gelling et al., 2003), *Gcgr*^*-/-*^ mice exhibited marked α-cell hyperplasia with the glucagon content being ∼10-fold higher than in wildtype islets and rate of glucagon release correspondingly increased (Figure 6C). For display purposes glucagon secretion in *Gcgr*^*-/-*^ islets was normalized to that at 1 mM glucose. Unlike what was observed in wildtype islets, neither GLP-1(7-36), nor GLP-1(9-36) exerted any statistically significant glucagonostatic effects in *Gcgr*^*-/-*^ islets (Figure 6B). It should be noted that that these islets likewise failed to respond to elevated glucose with suppression of glucagon secretion, suggesting that the function of α-cell in *Gcgr*^-/-^ mice may extend beyond the permanent loss of glucagon signaling. Notably, ablation of GCGR did not affect glucagon secretion at 1 mM glucose and in both wild-type and *Gcgr*^*-/-*^ islets the fractional rate of release was ∼0.1%/h (Figure 6D).

### Glucagon receptor antagonists abolish glucagonostatic effect of GLP-1(9-36)

To avoid long-term/compensatory effects of genetically ablating the GCGRs, we compared the glucagonostatic effects of physiological concentrations of GLP-1(9-36) in mouse islets in the absence and presence of the glucagon receptor antagonist L-168049 (Cascieri et al., 1999).

Under control conditions, GLP-1(9-36) (10 pM) inhibited glucagon secretion in mouse islets by ∼50% (Figure 6E). When tested in the presence of L-168049, GLP-1(9-36) failed to affect glucagon secretion. In the same islets, elevation of glucose to 6 mM inhibited glucagon secretion ∼50% (confirming that the α-cells were functional). These experiments were repeated in human islets. We observed that GLP-(9-36) (10 pM) inhibited glucagon secretion in human islets almost as strongly as 6 mM glucose (48% and 56%, respectively) and that the effect of the GLP-1 metabolite was abolished by the glucagon receptor antagonists desHis^1^Pro^4^Glu^9^glucagon (“Peptide N) and desHis^1^Pro^4^Glu^9^Lys^12^FA-glucagon (‘Peptide R) (O’Harte et al., 2014) (Figure 6F).

In both mouse and human islets, glucagon secretion at 1 mM glucose was not affected by inclusion of the glucagon receptor antagonists in the incubation media. Taken together with the observation in *Gcgr*^*-/*-^ islets it can be concluded that glucagon is not a feedback regulator of its own release.

### Effects of GLP-1(9-36) in vivo

Plasma glucose reflects the balance between the hypoglycemic action of insulin and hyperglycemic action of glucagon. The observation that GLP-1(9-36) inhibits glucagon secretion *in vitro* therefore suggests that it may affect systemic glucose homeostasis. GLP-1(9-36) was administered intraperitoneally in mice at a dose of 500 ng/g body weight as reported previously by other investigators (Day et al., 2017). Plasma GLP-1(9-36) peaked after 10 min, attaining a maximal concentration of 400 pM. The concentration then decayed exponentially but remained >60 pM (5-fold higher than basal) 60 min after injection of the peptide (Figure S7A), close to that producing the maximum glucagonostatic effect in the *in vitro* measurements (cf. Figure 1C) but it was without effect on plasma glucose and glucagon when administered to fed mice with a plasma glucose of 11±0.5 mM (n=7) (Figure S7B-C).

We reasoned that the systemic effects of GLP-1(9-36) might be more apparent when glucagon secretion is stimulated by insulin-induced hypoglycaemia. In mice fasted for 5h, insulin (0.5 U/kg body weight) lowered plasma glucose from a basal 7.5 mM to 4 mM (Figure 7A). This was associated with a ∼10-fold increase in plasma glucagon at 30 and 45 min after induction of hypoglycemia (Figure 7B). When insulin was co-injected with GLP-1(9-36), plasma glucose followed the same trajectory as when insulin was administered alone (Figure 7A) but the stimulation of glucagon secretion was reduced by ∼40% (Figure 7B). This inhibition is similar to that observed in isolated islets (Figure 1C-D). Following more extended fasting, a stronger hypoglycemic effect of GLP-1(9-36) was observed (Figure S7D-F).

**Figure 7:**
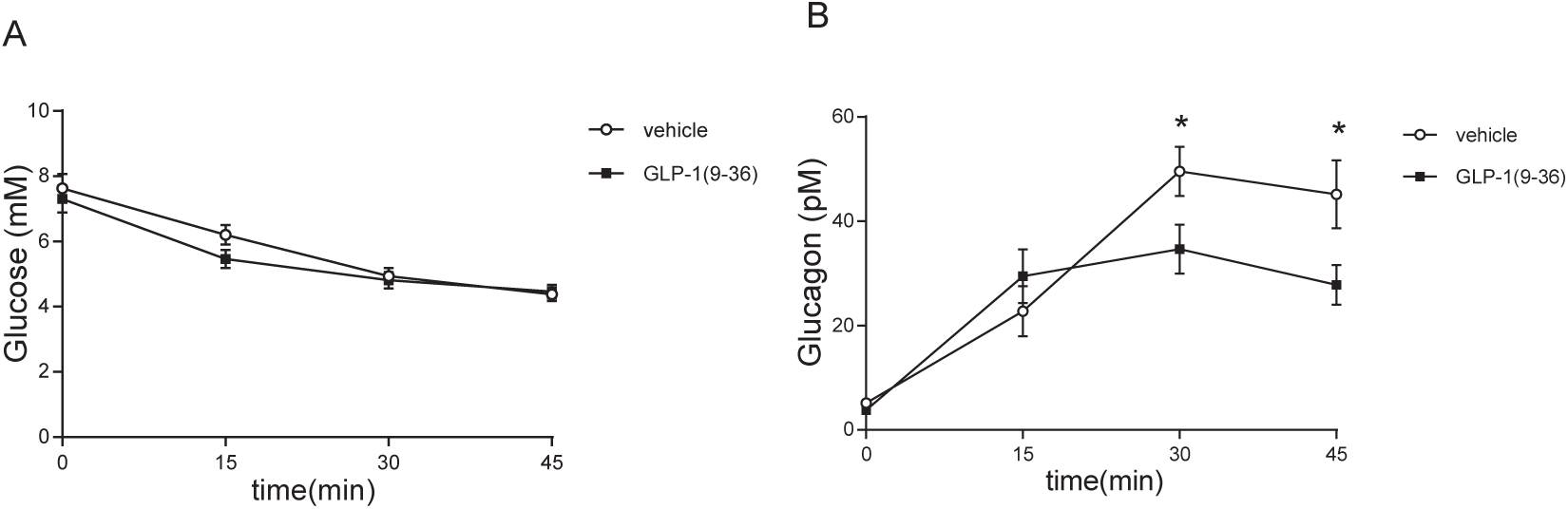
GLP-1(9-36) effects on glucose and glucagon secretion during insulin-induced hypoglycemia *in vivo*. (**A-B**) Effect of GLP-1(9-36) injection on glucose (A) and on glucagon (B) after 15, 30 and 45 minutes from insulin administration, in mice fasted for 5h (n=11-13) *P<0.05 for indicated comparisons.

## Discussion

Here we have studied the effects of GLP-1(7-36) and its metabolite, GLP-1(9-36) on pancreatic islet function. We propose that GLP-1(7-36) and GLP-1(9-36) inhibit glucagon secretion by a combination of (at least) two mechanisms that involve distinct receptors and intracellular signal transduction pathways. Whereas one of these is mediated by binding of GLP-1(7-36) to the cloned GLP-1R, the second mechanism is mediated by the degradation product GLP-1(9-36), involves binding to the glucagon receptor (GCGR) and culminates in activation of a pertussis toxin-sensitive inhibitory GTP-binding protein.

Whereas the two mechanisms normally cooperate, they may (under certain experimental conditions) individually suffice to maximally suppress glucagon secretion. Accordingly, genetic ablation of one receptor subtype may not be enough to produce a clear phenotype and combination of genetic and pharmacological strategies are therefore required to unveil the role of the different receptors. We emphasize that our findings are fully consistent with the previously proposed concept (De Marinis et al., 2010; Ramracheya et al., 2018) that GLP-1 exerts its glucagonostatic action by intrinsic effects in the glucagon-secreting α-cells that are independent of paracrine effects from the neighboring islet cells. Figure S8 outlines a model that explains our findings and that depicts the crosstalk between GLP-1R-dependent and –independent (GCGR) pathways in α-cells in the islet.

### GLP-1 signaling in α-cells

Plasma GLP-1 levels range between 10-50 pM of which only <20% is GLP-1(7-36) (Holst, 2007). In isolated islets, glucagon secretion was strongly inhibited by GLP-1(7-36) and (9-36) at concentrations as low as 1-10 pM (Figure 1). Accordingly, glucagon secretion is under strong tonic inhibition by circulating levels of GLP-1(7-36) and GLP-1(9-36). This may explain the observation that cell-specific ablation of *Glp1r* in α-cells leads to increased plasma glucagon in fed mice (Zhang et al., 2019).

The physiological role of the glucagonostatic effect of GLP-1(9-36) remains to be established but it may serve to restrict glucagon secretion following a mixed meal, for example, the increase in circulating amino acids would otherwise lead to strong stimulation of glucagon secretion, which would be functionally inappropriate.

Whether *Glp1r* is present in glucagon-secreting α-cell remains debated but most studies agree that it is expressed at much lower levels than in the neighboring β-cells (Blodgett et al., 2015; DiGruccio et al., 2016; Richards et al., 2014). In this study, expression of *Glp1r* in α-cells was only ∼2% of the non-α-cells (i.e. β- and δ-cells) (Figure S3). As expected, genetic ablation of *Glp1r* abolished the potentiating effect of GLP-1 on glucose-induced insulin secretion. However, the glucagonostatic effects of both GLP-1(7-36) and (9-36) were unaffected.

### GLP-1R- and PKA-dependent effects of GLP-1 in α-cells

Despite the low expression of *Glp1r* in α-cells and the finding that the glucagonostatic effect of GLP-1 was seemingly intact in *Glp1r*^*-/-*^ mice, it nevertheless appears that these receptors contribute to the glucagonostatic action of GLP-1(7-36). This is suggested by observations made with/without pretreatment with pertussis toxin (PTX). In wild-type islets, the glucagonostatic effect of GLP-1(7-36) is unaffected by pretreatment with PTX. In *Glp1r*^*-/-*^ mice, GLP-1(7-36) was still inhibitory under control conditions but failed to suppress glucagon secretion after PTX pretreatment.

How GLP-1(7-36) exerts its glucagonostatic effect remains unclear but we have previously proposed that activation of the few receptors present on the α-cells leads to a small increase in intracellular cAMP, sufficient to activate PKA with resultant inhibition of glucagon secretion (De Marinis et al., 2010). This also explains our present finding that the PKA-inhibitor 8-Br-Rp-cAMPS reduces the inhibitory effect of GLP-1(7-36) by 50% (Figure 3A). The downstream PKA-dependent mechanism leading to inhibition of glucagon secretion remains to be identified but there is evidence that the effect is mediated by inhibition of high voltage-activated P/Q-type Ca^2+^ channels sensitive to ω-agatoxin (Ramracheya et al., 2018) (Figure S8). This scenario is supported by the findings that low concentrations of the adenylate cyclase activator forskolin as well as adrenaline inhibit rather than stimulate glucagon secretion and it is only at high concentrations of the compounds that stimulation of glucagon secretion is observed (De Marinis et al., 2010). The latter effect may contribute to the observed paradox that high (nM) concentrations of GLP-1 are less inhibitory on glucagon release than low (pM) concentrations of the hormone; this may be because high concentrations of GLP-1(7-36) increase intracellular cAMP slightly more than required to produce maximal inhibition of glucagon release. We also considered the possibility that the weaker inhibitory effect may reflect GLP-1R receptor desensitization (35) but it is notable that 1 pM GLP-1(7-36) remained strongly inhibitory when applied after a 2-min exposure to 10 nM of the incretin; if receptor desensitization had occurred, then its inhibitory effect should have been reduced compared to when the low concentration was applied to begin with.

### Pertussis toxin-sensitive and GLP-1R-independent effects of GLP-1 in α-cells

In addition to the cAMP/PKA-dependent actions of GLP-1(7-36) mediated by activation of GLP-1R, GLP-1(9-36) also inhibits glucagon secretion by a pertussis toxin-sensitive (G_i_) effect, involving membrane receptors other than GLP-1R. The finding that GLP-1(7-36) is without effect when applied in the simultaneous presence of exendin(9-39) (to block the GLP-1Rs) and sitagliptin (to inhibit the activity of DPP-4s) (Figure 4E-F) further suggests that its inhibitory effect on glucagon is mediated following its degradation to GLP-1(9-36) (Figure S8B). These observations may help to explain the reported improved glycaemic control in patients treated with a combination of the GLP-1R agonist exenatide (exendin-4) and the DPP-4 inhibitor sitagliptin (Violante et al., 2012).

Receptor binding studies using the cloned human GCGR suggest that they are activated by GLP-1(9-36) and that this culminates in activation of G_i_. Indeed, GCGR antagonists prevent the glucagonostatic effect of GLP-1(9-36). The suppression of glucagon secretion produced by GLP-1(9-36), but not that by GLP-1(7-36), was prevented in islets pretreated with pertussis toxin. Activation of GCGR by GLP-1(9-36) was enhanced by RAMP2, which is expressed at much higher levels in human α-cells than human β-cells. This may explain why high circulating levels of GLP-1(9-36) do not activate G_i_ in β-cells with resultant (equivalent to functionally-inappropriate) suppression of insulin secretion. How GLP-1(9-36) and activation of G_i_ inhibit glucagon secretion remains to be documented but we have previously shown that agents that activate G_i_ (including somatostatin) inhibit glucagon granule exocytosis in both mouse (Gromada et al., 2001) and human (Kailey et al., 2012) α-cells by depriming of release-competent secretory granules (possibly via activation of the protein phosphatase calcineurin (Renstrom et al., 1996) and dephosphorylation of proteins involved in exocytosis such as SNAP-25 (Nagy et al., 2004)) and it is possible that GLP-1(9-36) may have the same effect. We acknowledge that expression of *Gcgr* is low in α-cells but a role of glucagon receptors is supported by the finding that GLP-1 had no statistically significant glucagonostatic effect in *Gcgr*^*-/-*^ islets.

It would appear counterintuitive that picomolar concentrations of GLP-1(9-36) inhibit glucagon secretion by activation of GCGRs and yet high endogenous glucagon (>30 nM; (Svendsen et al., 2018)) does not exert any glucagonostatic effects. One potential explanation could be that the kinetic rates for the binding of the agonists to the receptor differ. It is at least theoretically possible that when GLP-1(9-36) binds to the GCGR, it has a slow off rate resulting in prolonged signalling effects compared to glucagon. A second alternative could relate to the extent of agonist-induced internalisation of the receptors. While glucagon may induce rapid GCGR internalisation, GLP-1(9-36) may not, resulting in a sustained and prolonged response. We acknowledge that the EC_50_ for the stimulation of cAMP production in the receptor binding studies are >100-fold higher than the EC_50_ for the glucagonostatic effects observed in the secretion studies. However, it is possible that there is a strong ‘amplification’ of the response and that the downstream effector mechanisms saturate even when receptor activation is submaximal.

### Coda

*Beyond the* α*-cell.* These findings suggest that GLP-1 exerts, at least part of, its effects on glucagon secretion by a *Glp1r*-independent mechanism as also suggested for its cardioprotective effects in the rodent heart (Ban et al., 2010). There have been multiple reports that GLP-1(7-36) and GLP-1(9-36) exert effects on the CNS, cardiovascular system, gastrointestinal tract, liver and muscle that persist in the absence of the classical GLP-1 receptor and which are likely to involve distinct receptor(s)/mechanisms (Cantini et al., 2016; Davidson, 2011; Drucker, 2016; Ussher and Drucker, 2012). More recent evidence indicates that even GLP-1 breakdown products can exert beneficial physiological effects (50). The existence of an alternative GLP-1R has been inferred for a long time but attempts to identify it have failed and there is no obvious candidate in the genome (see http://www.glucagon.com/glp1secondreceptor.html). Our data suggest that an alternative GLP-1R does not exist and that *Glp1r*-independent effects of GLP-1(7-36) and GLP-1(9-36) instead reflect the simultaneous response to other (related) GPCRs (that normally mediate the effects of other ligands such as glucagon) and which can be modulated by the presence of tissue-specific accessory proteins such as RAMP2. Indeed, there is growing recognition that proglucagon-derived peptides bind promiscuously to their cognate receptors and that glucagon is a non-conventional GLP-1R agonist (Chepurny et al., 2019). It is possible that the concept of receptor ‘promiscuity’ can be extended to other cases where a hormone exerts an effect without an obvious candidate receptor.

## STAR ⍰ Methods

### Human islets

Human pancreatic islets were isolated (with ethical approval and clinical consent) at the Diabetes Research and Wellness Foundation Human Islet Isolation Facility (OCDEM, Oxford, UK) from the pancreases of at least 22 non-diabetic donors. Donors (10 females, 14 males) were on average 48 years old (range 19–61) with a BMI of 28 (range 21–37) (Table S1). Islets were isolated as previously described (Ramracheya et al., 2010). Human islets were usually released for experimental work within 24h of islet isolation. During the interval between islet isolation and the hormone secretion studies, islets were maintained in complete RPMI medium containing 5 mM glucose for up to 2 days prior to the experiments.

### Animals, islet isolation and whole pancreas perfusion

Most studies were conducted in 8-16 weeks old mice on a C57BL6/J or NMRI background.

### Ethical approval

All animal experiments were conducted in accordance with the ethical guidelines of the Universities of Oxford, Lausanne, Copenhagen and Goteborg and were approved by the local Ethics Committees.

Mice were housed at up to six per cage and kept on 12h light-dark cycle with free access to chow diet and water.

In addition to the ordinary mice, three different mouse strains were used in this study: global *Glp1r*^*-/-*^ mice (Scrocchi et al., 1996); global *Gcgr*^*-/-*^ mice (Gelling et al., 2003); and α-cell-specific *Gpr119*^*-/-*^ mice (GLU-Cre x GRP119(fx/fx) mice) (Moss et al., 2016).

For experiments with the mouse knockout models, sex- and age-matched wild type littermates were used as controls.

The mice were killed by cervical dislocation, the pancreas quickly resected and pancreatic islets isolated by liberase (Sigma) digestion. The islets were handpicked, size-matched and grouped manually.

### Reagents

GLP-1(7-36), GLP-1(9-36), exendin-4 and exendin(9-39) were purchased from Bachem (Weil am Rhein, Germany). The PKA inhibitor 8-Br-Rp-cAMPS (Rp-cAMPS) was purchased from BioLog Life Science Institute (Bremen, Germany), the SSTR2 antagonist CYN154806 from Tocris Bioscience (Bristol, UK), the Gpr119 agonist AS1269574 from Tocris and pertussis toxin from Sigma. All other reagents were of analytical grade. Human GLP-1(7-36)NH_2,_ GLP-1(9-36)NH_2_, glucagon (GCG) and exendin-4 (Ex-4) were custom synthesised by Generon (Slough, U.K.). GPR119 small molecule agonist, AR231453, was purchased from Sigma (Dorset, U.K.) and glucagon receptor (GCGR) small molecule antagonist, L-168049 was purchased from Tocris (Bristol, U.K.). Rolipram and forskolin were purchased from Cayman Chemical Company (Michigan, U.S.). All peptide ligands were made up to 1mM stock solution in deionised water while small molecule compounds, except for rolipram (which was made up to 25 mM), were made up to 10mM in dimethyl sulfoxide (DMSO). All compounds and peptide ligands were stored at −20°C before assays. cDNA constructs of the human GCGR was donated by Professor Patrick Sexton (Monash University, Australia) while cDNA constructs of human GPR119 were purchased from cDNA.org. The LANCE® cAMP detection assay kits and 384-well white Optiplates were purchased from PerkinElmer Life Sciences (Waltham, U.S.A.). Minimal Essential Medium (MEM) and heat inactivated fetal bovine serum (FBS) were purchased from Gibco™ (Thermo-fisher Scientific, U.K.).

### Measurements of islet hormone secretion

Mouse islets were used acutely, except for studies with PTX, where islets were treated with the toxin for at least 4 hours, and were, pending the experiments, maintained in tissue culture for <24 h in RPMI medium containing 10% fetal bovine serum (FBS), 1% penicillin/streptomycin and 10 mM glucose prior to the measurements. Human islets were maintained in culture for up to 48 h in RPMI medium containing 10% fetal bovine serum (FBS), 1% penicillin/streptomycin and 5 mM glucose.

Experiments were conducted using batches of 10–15 size-matched islets per tube (in triplicate) as previously described (Ramracheya et al., 2010; Vergari et al., 2019). We note that glucagon secretion exhibits variability between preparations. To circumvent these confounds, each donor/group of mice were used as its own control when testing the effect of a compound.

Islets were washed twice in RPMI prior to preincubation in Krebs-Ringer buffer (KRB) containing 2 mg/ml BSA (S6003, Sigma-Aldrich) and 3 mM glucose for 1 h at 37°C. Following this, islets were incubated in 0.3 ml KRB with 2 mg/ml BSA, supplemented with various glucose concentrations or compounds as indicated. After incubation, the supernatant was removed and quickly frozen and stored at −80°C. The islet pellets were lysed with 0.1 ml ice-cold acid ethanol. Insulin and glucagon were determined by radioimmunoassay (Millipore and Oxford Biosystem, respectively) according to the manufacturers’ instructions. The concentration of glucagon secreted by the islets ranged between ∼1.5-7.5 pg/islet/h (∼100-450 pg/ml) for mouse islets and between ∼2-10 pg/islet/h (∼140-590 pg/ml) for human islets. Whole pancreas perfusion studies were done as described previously (51).

### Receptor binding studies

#### Cell Culture and transient transfection

HEK 293 cells lacking the calcitonin receptor (ΔCTR-HEK 293 cells) were given by Drs. David Hornigold, Jacqueline Naylor and Alessandra Rossi (MedImmune, Cambridge, UK), and used as described elsewhere (Bailey et al., 2019). ΔCTR-HEK 293 cells were cultured in MEM supplemented with 10% heat-inactivated FBS plus 1% non-essential amino acids. The cells were incubated in a humidified 95% air/5% CO_2_ incubator at 37°C and were used between passages 1 to 5. ΔCTR-HEK 293 cells were transfected with polyethylenimine (PEI) as per manufacturer’s protocol using 1:6 (w:v) DNA:PEI ratio on 24-well plate. The transfected cells were grown 48 h prior to assays.

#### cAMP accumulation assays

Assays were performed as previously described (Knight et al., 2016; Weston et al., 2015; Weston et al., 2014). In brief, ΔCTR-HEK 293 cells transiently transfected with GCGR or GPR119 were washed with phosphate buffered saline (PBS), resuspended in PBS containing 0.1% bovine serum albumin (BSA) plus 25*µ*M rolipram, and seeded at 2000 cells/well in 384-well white Optiplates. Agonists were serially diluted at concentrations in the range of between 100 μM to 0.01□pM (depending upon agonist used) in a 96 well-plate. GCGR antagonist L-168049 was added to the serially diluted peptide ligands in the antagonist assay and the DMSO content was kept at 2% across all wells. The cells were stimulated with ligands for 15 or 30 min as indicated and cAMP accumulation was measured using a LANCE® cAMP detection assay kit. Plates were read using a Mithras LB 940 multimode micro-plate reader (Berthold Technologies). Values were converted to concentration using a cAMP standard curve performed in parallel and standardised to cells stimulated with 100 μM forskolin, which provided the system maximum.

### Plasma glucose measurements during insulin tolerance tests

Fed blood glucose levels (data point before fasting) were measured from a blood drop obtained by a tail vein nick using the Accu-Chek Aviva kit (Roche Diagnostic). The mice were then fasted for 5h prior to the experiments. Fast-acting human insulin (Actrapid, Novo Nordisk) was injected intraperitoneally at the indicated doses with a 25-gauge needle at time zero. GLP-1(9-36) was injected intraperitoneally with insulin as indicated. In the control experiments, insulin was co-injected with the solvent.

Tail vein blood glucose and glucagon levels were monitored using a glucometer before and an ELISA (Mercodia Glucagon ELISA, Uppsala, Sweden). Total GLP-1 was measured by ELISA (Crystal Chem, Zaandam, Netherlands).

### Statistics

All data are reported as mean ± SEM, unless otherwise stated. Statistical significance was defined as P<0.05. All statistical tests were conducted in Prism 5 (GraphPad Software, San Diego, CA). For two groupings, a t-test was conducted. If the data were nonparametric, a Mann–Whitney test was conducted. For more than two groupings, a one-way ANOVA was conducted. If there were two independent variables, a two-way ANOVA was conducted. For secretion data, a minimum of two human donors were used and each replicate was considered an individual experiment. For experiments on mouse islets, each replicate (using different groups of islets) was regarded as a separate experiment and the number of mice used for each experiment is stated and, for each experiment, at least 4 mice were used. Because data were collected over several years, with mice of different ages and that it was not possible to test all conditions with the same batch of islets, glucagon secretion has in most Figures been expressed after normalization to basal measured at 1 mM glucose (unless otherwise indicated). However, basal glucagon secretion values are reported for all panels.

### Study approval

All experiments on mouse islets were conducted in accordance with the United Kingdom Animals (Scientific Procedures) Act (1986) and the University of Oxford ethical guidelines. Breeding of *Gcgr* null mice were in compliance with an animal experiment license issued by the Danish Committee for Animal Research and approved by the local animal welfare committee at the University of Copenhagen. Breeding of *Glp1r*-deficient mice was performed at the University of Lausanne and were reviewed and approved by the Veterinary Office of Canton de Vaud.

Human pancreases were obtained with ethical approval of the NHS National Research Ethics Centre, Oxfordshire, United Kingdom (REC B) and clinical consent from heart-beating donors.

## Acknowledgements

We thank Prof Paul Johnson and his team at the University of Oxford for access to donor human islets. We also thank Prof Jens Juul Holst and Dr Berit Svendsen at the University of Copenhagen for provision of the GCG-R knockout mice. We thank Dr F O’Harte (University of Ulster, Coleraine, Northern Ireland, UK) for the kind gift of the glucagon receptor antagonists Peptide N and R and Professor Niels Billestrup at the University of Copenhagen for access to his laboratory for the experiments on the GCGR mice. We also thank Mr David Wiggins, Mr Samuel Stephen and Miss Natasha Allen for technical assistance.

The work was supported by a Diabetes UK RD Lawrence Fellowship to Dr Reshma Ramracheya, a Novo Nordisk University of Oxford postdoctoral fellowship to Dr Claudia Guida and an OXION Wellcome Trust studentship to Miss Joely Kellard.

Work in Oxford was also supported by a Wellcome Trust Senior Investigator Award held by Prof Patrik Rorsman.

Studies in Göteborg were covered by a Wallenberg Scholars Fellowship from the Knut and Alice Wallenbergs Stiftelse and an International Recruitment Award from the Swedish Research Council (VR) to Patrik Rorsman and a VR Research Grant to Ingrid Wernstedt Asterholm.

Studies in Cambridge were supported by the Rosetrees foundation (to H. Y. Y and G.L), a Wellcome Trust joint investigator award (FR) an MRC programme within the Metabolic Diseases Unit (FR). H.Y.Y was supported by an international scholarship from the Cambridge Trust. Work in Lausanne (BT) was supported by grants to B.T. from the Swiss National Science Foundation and the European Research Council.

## Author contributions

RR, CG, CM, IWA, DB, AB, BC, MVC, MH, JK, LJM, JR, NJGR, MS and HYY performed the experimental studies. Animal models were provided by FR and BT. GL directed the receptor binding studies. The study was conceived by PR and RR who planned and designed the experiments. PR wrote the manuscript. CG and RR coordinated the editing of the final version of the manuscript. All co-authors contributed to the discussion and approved the final version of the manuscript.

