## supplemental material and figures for "GLP-1(9-36) mediates the glucagonostatic effect of GLP-1 by promiscuous activation of the glucagon receptor"

*GLP-1 does not inhibit glucagon secretion by paracrine effects*

The capacity of GLP-1(9-36) to inhibit glucagon secretion was not associated with any stimulation of insulin secretion. Increasing glucose from 1 to 6 mM stimulated 2-fold increase in insulin secretion in mouse and human islets (Figure S2A-B). There were no stimulatory effects of GLP-1(9-36) (10 pM) at 6 mM glucose. We also confirmed that GLP-1(9-36) was ineffective at influencing insulin release when applied at 1 mM glucose (not shown). The failure of GLP-1(9-36) to stimulate glucose-induced insulin secretion in mouse islets is in agreement with previous reports and distinguishes it from the strong insulinotropic effect of GLP-1(7-36)(Vahl et al., 2003).

We also explored whether GLP-1(9-36) could affect glucagon secretion by a paracrine effect mediated by stimulation of somatostatin secretion from the delta-cells in islets. When applied in the presence of 1 mM glucose, GLP-1(9-36; 10 pM) did not affect somatostatin secretion in human pancreatic islets (Fig. S1C). Furthermore, the somatostatin receptor (SSTR) inhibitor CYN154806 (100 nM) did not affect the ability of GLP-1(9-36) to reduce glucagon secretion in mouse islets (Fig. S2D). Notably, the SSTR antagonist strongly increased glucagon secretion both in the absence and presence of GLP-1(9-36), consistent with the idea that glucagon release is under strong tonic inhibition by endogenously released somatostatin(Hauge-Evans et al., 2009). However, the relative inhibitory effect produced by GLP-1(9-46) remained ~50%.

*Role of Gpr119.* We reasoned that GLP-1 might suppress glucagon secretion by activation of GPR119(Schirra et al., 1998). *Gpr119* is expressed in mouse α-cells at levels comparable to *Adrb1* and *Gipr* (encoding the β-adrenergic type 1 and gastric inhibitory polypeptide (GIP) receptors, respectively) and at >10-fold higher levels than

*Glp1r* (the latter encoding the glucagon receptor) (Adriaenssens et al., 2016; Blodgett et al., 2015; DiGruccio et al., 2016).

We measured glucagon secretion in mouse pancreatic islets exposed to 1 mM glucose and increasing concentrations of the GPR119 agonist AS12695704)(Yoshida et al., 2010). Maximum inhibition occurred at 5-50nM with higher concentrations (≥500 nM) being ineffective (Figure S4A). The observed U-shaped dose-response curve resembles that which we have previously reported for increasing concentrations of adrenaline(De Marinis et al., 2010).

In wild type islets, AS1269574 inhibited glucagon, an effect that was abolished in islets from *Gpr119*$^{-/-}$ mice(Lan et al., 2009) (Figure S4B-C) confirming the specificity of the agonist for Gpr119. By contrast, the capacity of GLP-1(7-36) and GLP-1(9-36) to inhibit glucagon secretion was unaffected by genetic ablation of *Gpr119* (Figure S4B-C), ruling out the involvement of this receptor as mediator of the *Glp1r*-independent effects of GLP-1(7-36) and GLP-1(9-36).

*GLP-1 does not activate GPR119 or GIPR*

Human GPR119 receptors expressed in HEK 293 cells(Bailey et al., 2019) did not respond to GLP-1(7-36) or GLP-1(9-36) with increased cAMP accumulation (Figure S4D). By contrast, The GPR119-selective agonist AR231453 stimulated cAMP in these cells.

In addition, we have also excluded the possibility that the GLP-1R agonists mediate their actions through the activation of GIPRs (Figure S5).

*Effects of GLP-1(9-36)* in vivo

Plasma glucose reflects the balance between the hypoglycemic action of insulin and hyperglycemic action of glucagon. The observation that GLP-1(9-36) inhibits glucagon secretion *in vitro* therefore suggests that it may affect systemic glucose homeostasis. GLP-1(9-36) was administered at a dose of 500 ng/g body weight. Plasma GLP-1(9-36) peaked after 10 min, attaining a maximal concentration of 400 pM. The concentration then decayed exponentially but remained >60 pM (5-fold higher than basal) 60 min after injection of the peptide (Figure S7A). When applied on its own and in fed mice, GLP-1(9-36) neither affected plasma glucose, nor plasma glucagon (Figure S7B-C).

*Schematic illustrating the regulation of glucagon secretion by GLP-1 via promiscuous receptor activation.*

Figure S8 (A) Normally, GLP-1 exerts a glucagonostatic effect by a combination of two effects (1 and 2): 1) Activation of GLP-1 receptor (GLP-1R) activates a stimulatory GTP-binding protein (Gs) with resultant stimulation of adenylate cyclase (AC), elevation of intracellular cAMP and activation of protein kinase A (PKA), that consists of regulatory (R) and catalytic subunits (C). Via protein phosphorylation, this leads to inhibition of voltage-gated P/Q-type $Ca^{2+}$ channels, the activation of which normally triggers exoycotis of glucagon granules (SG), and resultant suppression of glucagon release. This pathway is inhibited the PKA-inhibitor Rp-cAMPS. In addition, GLP-1 (following its degradation to GLP-1(9-36), a reaction catalysed by dipeptidyl peptidase 4 (DPP4)]), activates the glucagon receptor (GCGR). This pathway leads to activation of an inhibitor GTP-binding protein (Gi), possibly by activation of calcineurin, and culminates in dephosphorylation of exocytosis-regulating proteins, leading to depriming of secretory granules and reduced glucagon exocytosis. This pathway is inhibited by the DPP4 inhibitor sitagliptin (STG) and pertussis toxin (PTX). (B) After genetic (ablation of *Glp1r*) or pharmacological (Exendin(9-39)) disruption of the GLP-1R, GLP-1(7-36) lacks its receptor (indicated by X) and accordingly only mechanism (2) operates. (C) After genetic (ablation of *Gcgr*) or pharmacological (e.g. L-168049) manipulation of the GCGR, GLP-1(9-36) lacks the receptor (indicated by X) but GLP-1(7-36) may still inhibit glucagon secretion by mechanism (1). The model explains why the effects of GLP-1(7-36) and (9-36) on glucagon secretion differ with regard to sensitivity to receptor blockade and pharmacological inhibition of the intracellular signal transduction pathways.

### Supplemental Methods

*Flow cytometry of islet cells (FACS).* Pancreatic islets from either proglucagon- RFP were isolated as described above and single cells were isolated by trypsin digestion and mechanical dissociation as described previously(Vergari et al., 2019).

Single cells were passed through a MoFlo Legacy (Beckman Coulter). Fractions of RFP-positive or –negative cells were isolated by combining several narrow gates. Forward and side scatter were used to isolate small cells and to exclude cell debris. Cells were then gated on pulse width to exclude doublets or triplets. RFP-positive cells were excited with a 488 nm laser and the fluorescent signal was detected through a 580/30 nm bandpass filter. RFP-negative cells were collected in parallel.

*RNA extraction and quantitative RT-PCR.* Gene expression was analyzed by quantitative RT-PCR in a- (RFP+) and non-a-cell fractions (RFP-). Total RNA was isolated using a combination of TRIzol and PureLink RNA Mini Kit (Ambion, Thermofisher Scientific). On column DNase treatment was included to eliminate DNA contamination. cDNA was synthesized from 500 ng of total RNA using the High Capacity RNA-to-cDNA kit (Applied Biosystems, Thermofisher Scientific). Real time qPCR was performed using SYBR Green detection and gene specific QuantiTect Primer Assays (Qiagen) on a 7900HT Applied Biosystems analyser. All reactions were run in triplicates. Relative expression was calculated using ΔCt method, with GAPDH and PPIA used as reference genes.

*Fluorescent GIPR Ligand Binding.* Binding of GLP-1R agonists to the GIPR was assayed using a modified version of the Cisbio Tag-Lite® ligand binding assay (Schiele et al., 2015). HEK-293T cells were cultured for 24 hours in 6 well plates prior to transient transfection with pcDNA3.1-NLuc. Cells were subsequently grown overnight, harvested and seeded at 50000 cells/well into poly-L-lysine coated 96-well plates. The following day, growth media was removed, cells washed with PBS prior to the addition of 80 µl PBS containing 0.9 mM $CaCl_2.2H_2O$, 0.49 mM $MgCl_2.6H_2O$, and 0.1% BSA to each well. 10 µl of Nano-Glo® Substrate (Promega - diluted in PBS containing 0.49 mM $MgCl_2.6H_2O$, 0.9 mM $CaCl_2.2H_2O$ and 0.1% BSA) was added to each well to a final concentration of 0.1 µM and the plate was incubated for 5 min in the dark. 10 nM Tag-lite® GIPR Red Agonist (Cisbio) was added and the plate left to equilibrate for 5 min at room temperature. Emission was measured at 485 nm and 530 nm every 30 seconds for 2 minutes prior to the addition of “cold” 1 µM test compound (GIP (1-42), GLP-1 (7-36), GLP-1 (9-36), exendin-4, exendin 9-39) which was added by injection into each well to displace the bound Tag-lite® GIPR Red Agonist. Again, with emission was measured every 30 seconds for 10 min. Vehicle was added to each sample to represent a background control level of emission. The BRET signal was calculated by subtracting the 530 nm/485 nm emission ratio for vehicle treated cells from Tag-lite® GIPR Red Agonist treated cells. To correct for injection, changes in BRET (ΔBRET) for vehicle alone was subtracted from all data.

**Figure S1: GLP-1(7-36) inhibits glucagon secretion at low glucose concentration**

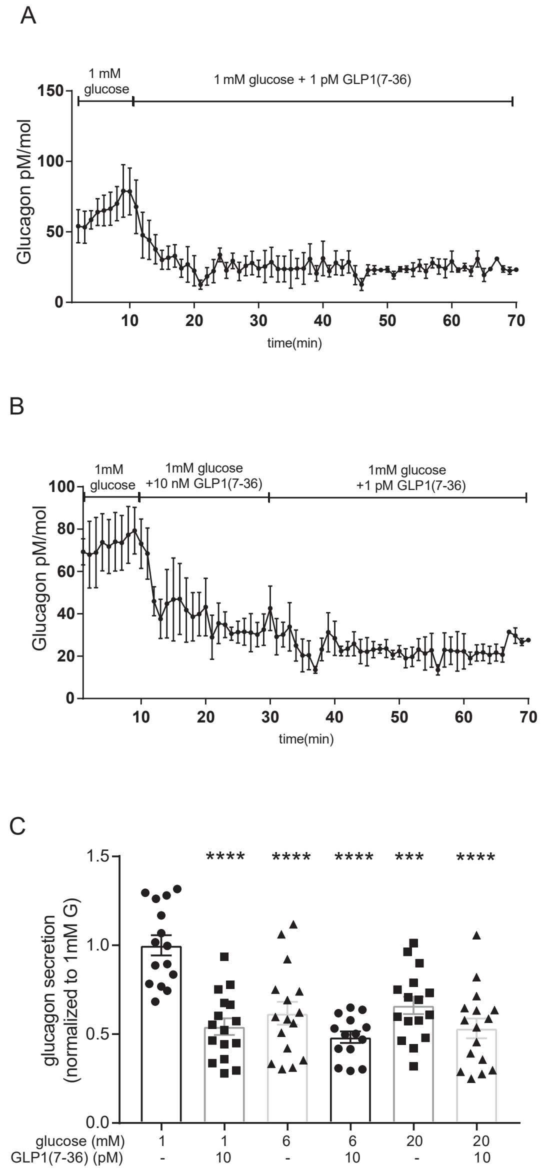


Figure S1 related to Figure 1. (**A-B**) Glucagon secretion in mouse pancreas perfused with 1mM glucose in presence of 1pM GLP-1(7-36) for 60 minutes (A) (n=3) or in presence of 10nM GLP-1(7-36) for 10 minutes and 10pM GLP-1(7-36) for other 40 minutes (B) (n=3).(**C**) Effect of GLP-1(7-36) on glucagon secretion in human islets at differentglucose concentration (n=15-16 using islets from 4 donors).**P<0.001, ****P<0.0001 versus 1mM glucose;1-way ANOVA with Dunnett's post-hoc test.

**Figure S2: GLP-1(9-36) does not modulate insulin and somatostatin secretion**

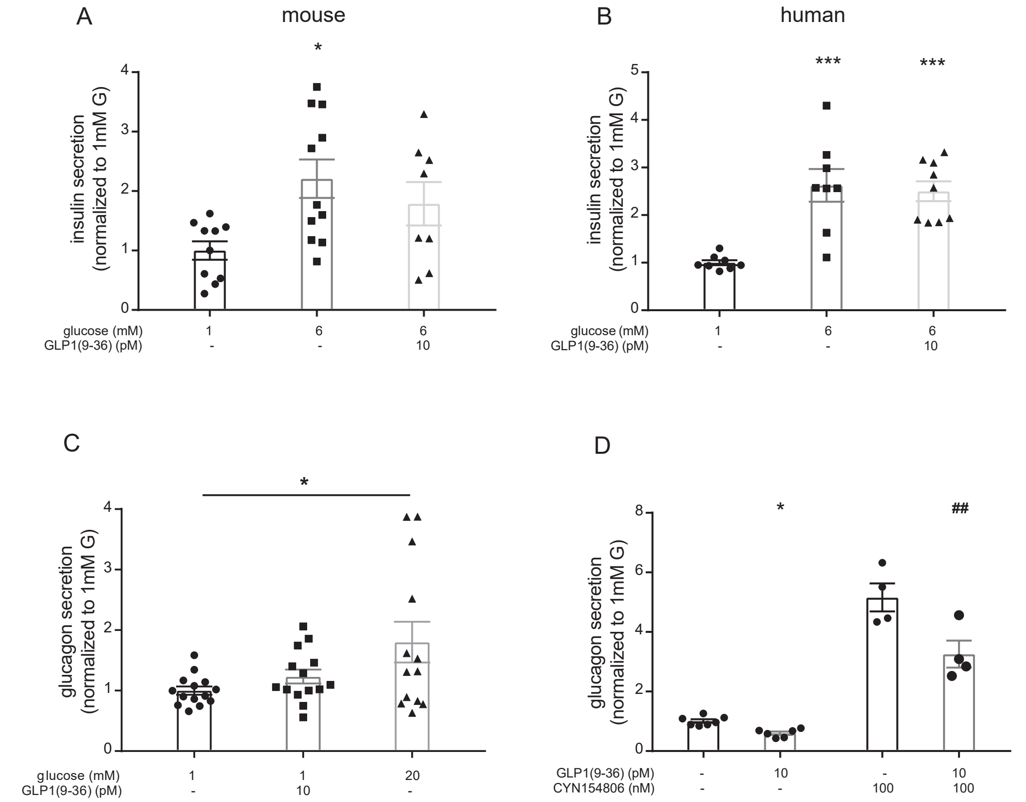


Figure S2 related to Figure 1 (**A-B**) Effects of GLP-1(9-36) on insulin secretion at 1 and 6mM glucose in mouse (A; n=8-11 using islets from 7 mice) and human (B; n=8 using islets from 3 donors) pancreatic islets. (**C**) Effect of GLP-1(9-36) on somatostatin secretion in human pancreatic islets (n=13 using islets from 4 donors) (**D**) Effects of GLP-1(9-36) on glucagon secretion in isolated mouse pancreatic islets in absence and presence of the SST receptor 2 inhibitor CYN154806 (n=4-6 using islets from 5 mice). *P<0.05, **P<0.01, ***P<0.001 versus 1 mM glucose; ## P<0.01 versus 1mM glucose+CYN154806, 1-way ANOVA with Dunnett's post-hoc test.

**Figure S3: Receptors expression in beta and alpha islet cells**

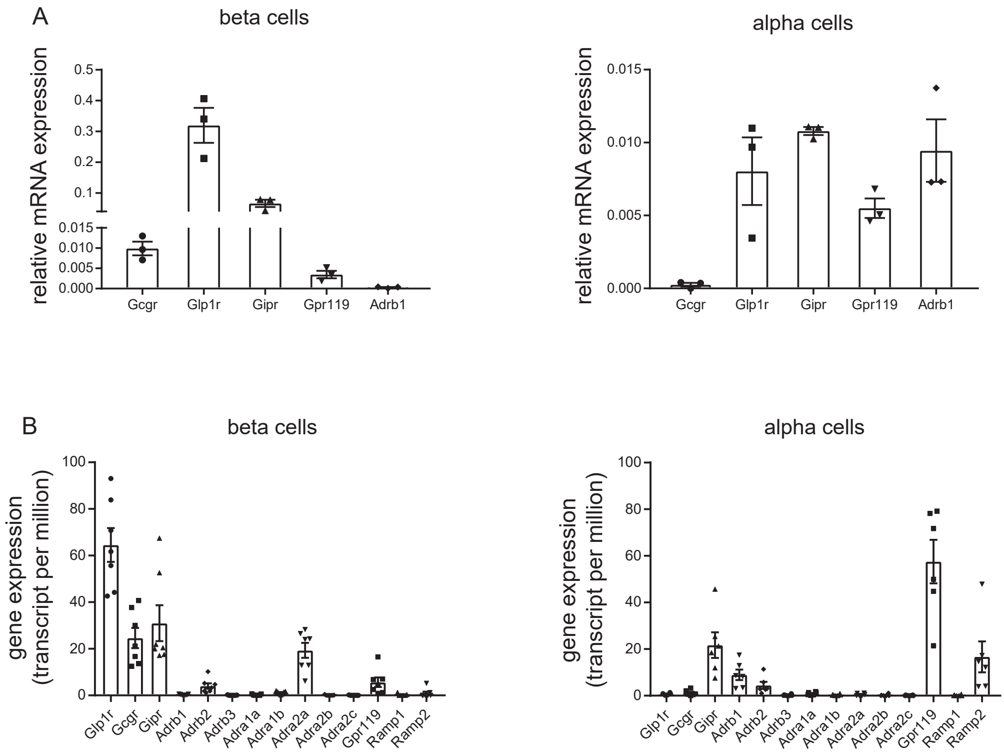


Figure S3. (**A**) relative mRNA expression of indicated genes in proglucagon- RFP -positive (alpha) or –negative (beta) murine islet cells (n=3) (**B**) RNA transcripts expression of indicated genes in beta (n=6) and alpha (n=5) human islet single cells as reported in Blodgett DM et al, 2015.

**Figure S4: Glp1r-independent effects of GLP-1 are not mediated by Gpr119**

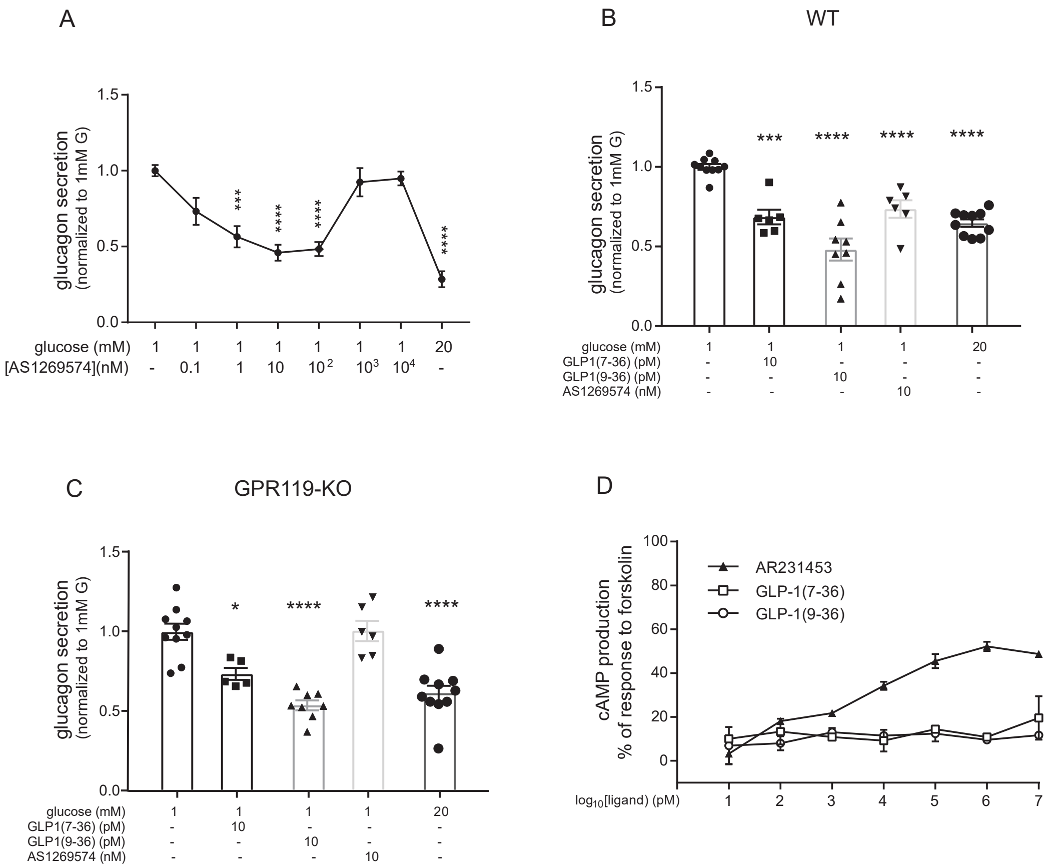


Figure S4. (**A**) Effects of increasing concentrations of the Gpr119 agonist AS1269574 on glucagon secretion measured in wild-type islets mice (n=6 using islets from 6 mice). (**B-C**) Effects of glucose, 10 pM GLP-1(7-36), 10 pM GLP-1(9-36) and 10 nM AS1269574 (included in the extracellular medium as indicated) on glucagon secretion in islets isolated from wild type (WT) and Gpr119-/- (GPR119-KO) mice (n=6-10 using islets from 8 mice). *P<0.05, ***P<0.001, ****P<0.0001 versus 1mM glucose; 1-way ANOVA with Dunnett's post-hoc test. (**D**) [ΔCTR] HEK 293 cells transiently transfected with GPR119 were stimulated for 30 mins with various potential agonists (n=6) and cAMP accumulation determined. Only the selective agonist AR231453 was able to induce a dose-dependent increase in cAMP accumulation. Data are expressed as percentage of cAMP production, determined using 100 μM forskolin stimulation.

### Figure S5: GLP-1(7-36) and GLP-1(9-36) do not activate Gipr

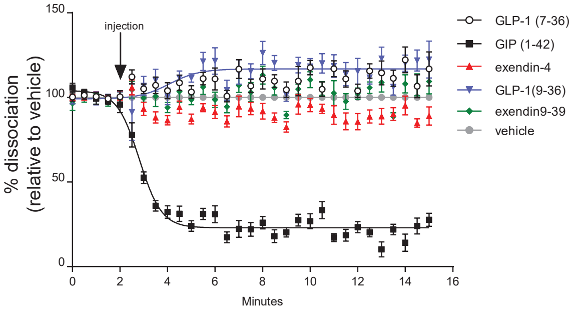


Figure S5. HEK-293T cells were transfected with NLuc-GIPR and displacement of 10 nM Tag-lite® GIPR Red Agonist was measured following injection of 1 µM test compound (GIP (1-42), GLP-1 (7-36), GLP-1 (9-36), exendin-4, exendin (9-39) after 2 min. To correct for injection, changes in BRET for vehicle alone was subtracted from all data sets.

**Figure S6: GLP-1(9-36) can activate the inhibitory Gi GTP-binding proteins**

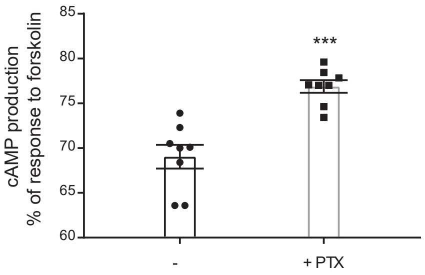


Figure S6. cAMP accumulation was determined when 10μM of GLP-1(9-36) was applied to [ΔCTR] HEK 293 cells transiently transfected with GCGR with or without 16 hour pre-treatment with PTX and stimulated for 15 min (n=8). ***P<0.001, for indicated comparisons; 1-way ANOVA with Dunnett's post-hoc test.

### Figure S7: Effects of GLP-1(9-36) *in vivo*

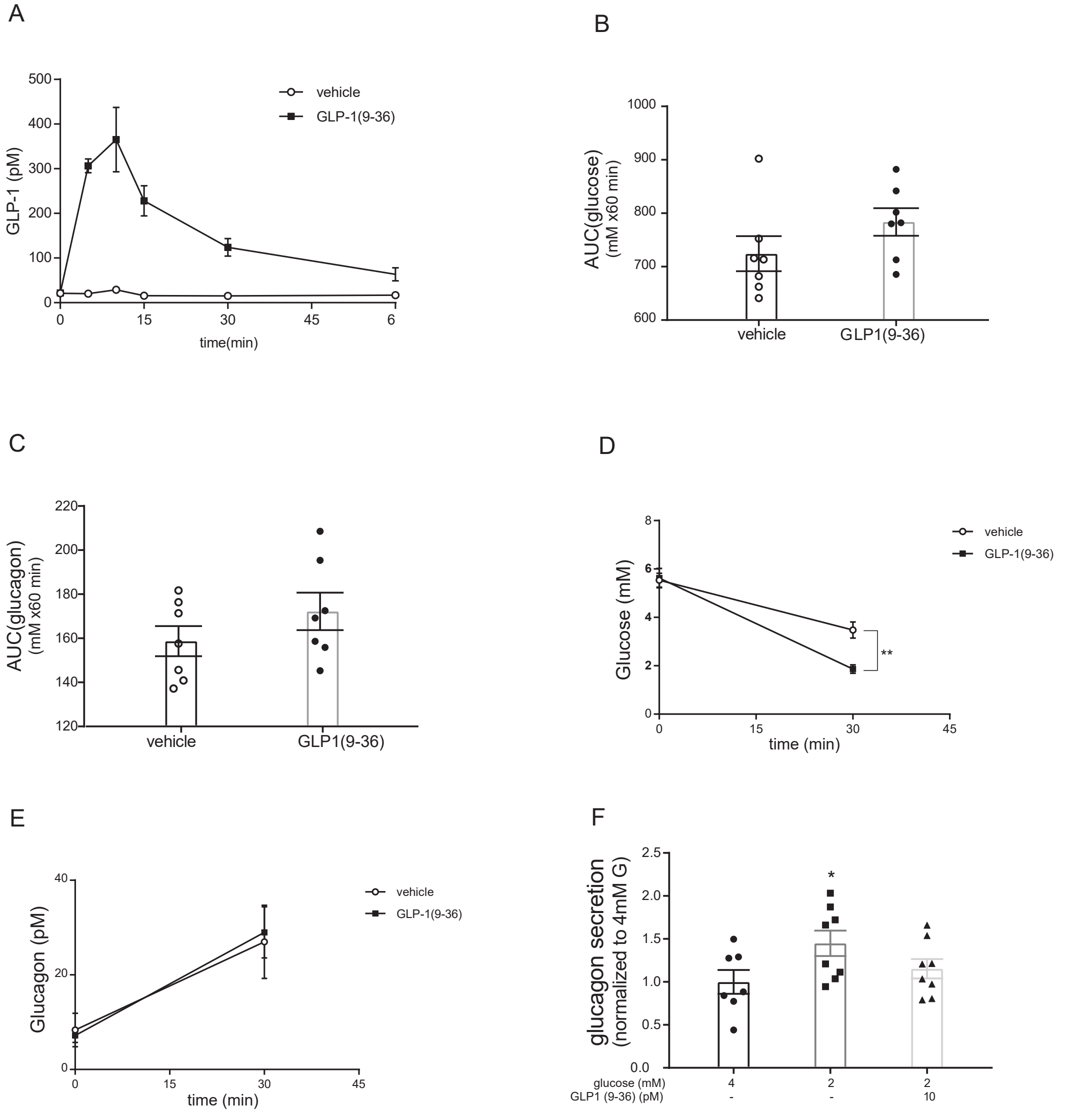


Figure S7 related to figure 7.(**A**) Plasma concentration of GLP-1(9-36) after injection of 500 ng/g body weight at t=0 in fed mice. (**B-C**) Plasma glucose (B) and glucagon (C) following injection of GLP-1(9-36). Data are presented as Area under the curve (AUC). (**D-E**) Effect of GLP-1(9-36) injection on glucose (D) and on glucagon (E) after 15,30 and 45 minutes from insulin administration, in mice fasted for 18h (n=5). **P<0.01 for indicated comparison.
(**F**) Effect of 10 pM GLP-1(9-36) on glucagon secretion in mouse islets (n=7-8 using islets from 4 mice, glucagon secretion at 2mM G=2.1 ±0.3 pg/islet) stimulated with 4mM and 2 mM glucose. *P<0.05 versus 4 mM glucose;1-way ANOVA with Dunnett's post-hoc test.

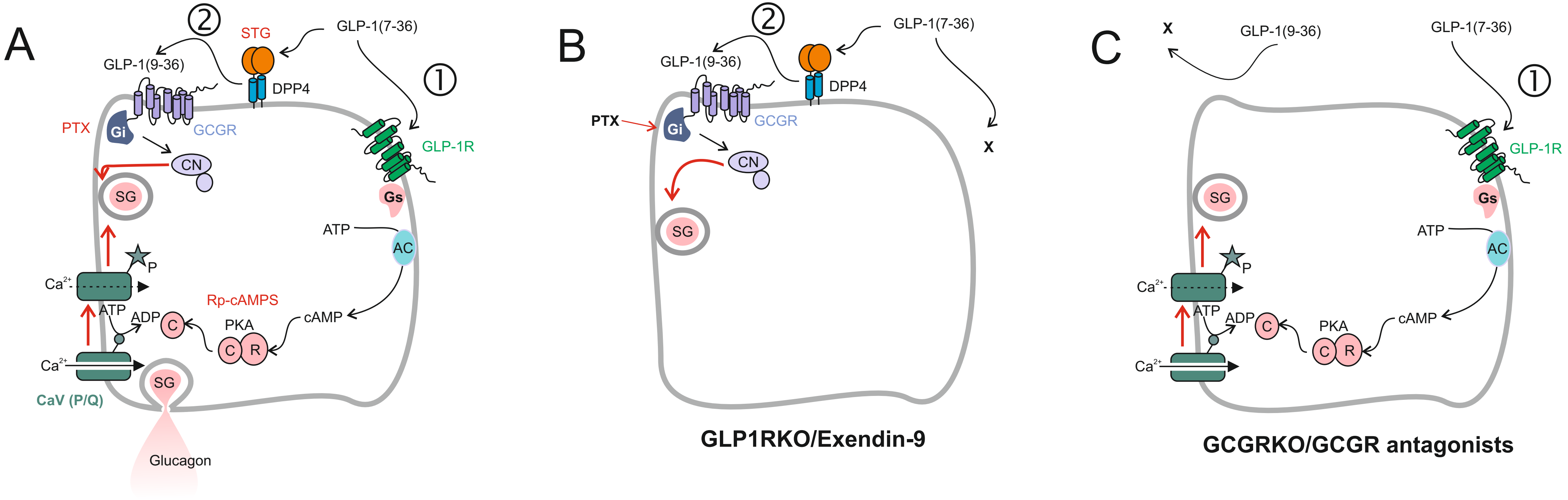
A
②
STG
GLP-1(7-36)
GLP-1(9-36)
DPP4
①
PTX
Gi
GCGR
GLP-1R
CN
SG
Gs
ATP
AC
P
Ca2+
Rp-cAMPS
cAMP
ATP
ADP
C
PKA
C
R
Ca2+
SG
CaV (P/Q)
Glucagon
B
②
GLP-1(7-36)
GLP-1(9-36)
DPP4
PTX
Gi
GCGR
X
CN
SG
GLP1RKO/Exendin-9
C
X
GLP-1(9-36)
GLP-1(7-36)
①
GLP-1R
SG
Gs
ATP
AC
P
Ca2+
ATP
ADP
C
cAMP
PKA
C
R
Ca2+
GCGRKO/GCGR antagonists

| **Table S1. Human islet donor details** | |
|---|---|
| **Age (yrs)** | 44.9 +/- 2.1 |
| **Male (n, %)** | 21, 75% |
| **BMI (kg/m$^2$)** | 28 +/- 0.9 |
| | Mean +/- SEM |
